# A Genetic Screen in *Drosophila* uncovers a role for *senseless-2* in surface glia in the peripheral nervous system to regulate CNS morphology

**DOI:** 10.1101/2024.01.09.574922

**Authors:** Haluk Lacin, Yuqing Zhu, Jose T. DiPaola, Beth A. Wilson, Yi Zhu, James B. Skeath

## Abstract

Despite increasing in mass approximately 100-fold during larval life, the *Drosophila* CNS maintains its characteristic form during this rapid growth phase. Dynamic interactions between the overlying basement membrane and underlying surface glia are known to regulate CNS structure in *Drosophila*, but the genes and pathways that establish and maintain CNS morphology during development remain poorly characterized. To identify genes that regulate CNS shape in *Drosophila*, we conducted an EMS-based, forward genetic screen of the second chromosome, uncovered 50 mutations that disrupt CNS structure, and mapped these alleles to 17 genes. Whole genome sequencing revealed the affected gene for all but one mutation. Identified genes include well characterized regulators of tissue shape, like *LanB1, viking, and Collagen type IV alpha1*, as well as previously characterized genes, such as *Toll-2* and *Rme-8*, with no known role in regulating CNS structure. We also uncovered that *papilin* and *C1GalTA* likely act in the same pathway to regulate CNS structure and found that the fly homolog of a glucuronosyltransferase, B4GAT1/LARGE1, that regulates Dystroglycan function in mammals is required to maintain CNS shape in *Drosophila*. Finally, we show that the *senseless-2* transcription factor is expressed and functions specifically in surface glia found on peripheral nerves but not those on the CNS proper to govern CNS structure, identifying a gene that functionally subdivides a glial subtype along the peripheral-central axis. Future work on these genes should clarify the genetic mechanisms that ensure the homeostasis of CNS shape and form during development.

## Introduction

Structure determines function is a unifying theme of biology. A thorough understanding of the genes that govern tissue or organ structure is, however, lacking. We use the *Drosophila* larval CNS as a model system to investigate the genetic factors that control tissue shape (Skeath, Wilson et al. 2017). The *Drosophila* larval CNS adopts its distinctive shape at the end of embryogenesis upon the shortening and thickening of the nerve cord in response to the concerted action of hemocytes, neural activity, and glial cell function in a process called nerve cord condensation (Olofsson and Page 2005, Karkali, Tiwari et al. 2022). Once its shape is established, the *Drosophila* CNS maintains its form throughout its massive growth during larval development.

The *Drosophila* larval CNS is consecutively and fully enwrapped by the CNS basement membrane and the perineurial and subperineurial surface glial cells, with the surface glia functioning as the blood-brain-barrier (Stork, Engelen et al. 2008, Meyer, Schmidt and Klambt 2014, Yildirim, Petri et al. 2019). The basement membrane is composed of independent Collagen IV and Laminin networks that are connected to each other by Perlecan; both networks establish connections to surface glial cells through interactions with the Integrin and Dystroglycan cell surface receptors (Yurchenco 2011). The tethering of the basement membrane to surface glial cells provides the CNS with structural and biomechanical strength essential to maintain its distinct shape. Interactions between the basement membrane and surface glia are, however, far from static. The tremendous growth of the CNS during larval life demands a dynamic basement membrane-surface glia interface that imparts structure to the CNS, while being continually remodeled to allow tissue growth and expansion.

In the larvae, the fat body secretes basement membrane proteins that are continuously deposited on the CNS basement membrane during development (Pastor-Pareja and Xu 2011), coordinating basement membrane deposition with tissue growth in the CNS. Remodeling of the basement membrane occurs at least in part through the action of the membrane-tethered and secreted extracellular proteases MMP1, MMP2, and AdamTS-A, which are expressed by surface glia (Page-McCaw, Serano et al. 2003, Meyer, Schmidt and Klambt 2014, Skeath, Wilson et al. 2017). These proteases oppose the tissue stiffening action of Collagen IV by cleaving basement membrane proteins and in so doing help maintain a delicate balance between tissue stiffness and softness that maintains tissue structure while allowing tissue growth and expansion (Apte 2009, Miller, Liu et al. 2011, Rodriguez-Manzaneque, Fernandez-Rodriguez et al. 2015, Skeath, Wilson et al. 2017). In addition to the fat body, migratory hemocytes also express basement membrane proteins in larvae and may serve as a second source of basement membrane proteins and a remodeling agent for the basement membrane (Isabella and Horne-Badovinac 2015).

To gain more insight into the factors that regulate CNS shape, we undertook a large-scale forward genetic screen of the second chromosome to identify mutations that disrupt CNS structure. Our screen identified 17 genes, including characterized regulators of tissue structure, characterized genes with no known role in controlling tissue structure, such as *Toll-2* (*18-wheeler*) *and Rme-8*, and previously uncharacterized genes, like a glucuronosyltransferase homologous to B4GAT1 and LARGE1/2, which regulate Dystroglycan in mammals, and the transcription factor Senseless-2. Our results highlight an unexpected role for perineurial glia in the peripheral nervous system (PNS) to help to establish and maintain CNS structure.

## Results and Discussion

The cellular and molecular processes that control CNS shape are poorly understood. To gain insight into these processes, we performed a large-scale EMS-based, forward genetic screen of the second chromosome to identify mutations that disrupt the morphology of the larval CNS (Fig. 1). Briefly, we used a target chromosome that contains a *3xP3-RFP* transgene insert in the second chromosome that labels most glia (Zhu, Cho et al. 2024), enabling rapid visualization of the CNS in live larvae. We set up 17,790 single male F1 crosses, screened 12,211 F3 lines for second chromosomal mutations that when homozygous were larval-, pupal-, or semi-lethal, identified 3180 such lines, and visually screened each for defects in overall CNS morphology. We identified 50 mutations that disrupt CNS shape, including 11 mutations in *LanB1*, *viking* or *Col4a1,* genes which have well characterized roles in regulating tissue and CNS structure (Pastor-Pareja and Xu 2011, Kim, Jeibmann et al. 2014, Skeath, Wilson et al. 2017, Dai, Estrada et al. 2018). The 39 remaining mutations grouped into three phenotypic classes: those with a wider CNS (n=1), a herniated or misshapen CNS (n=15), and an elongated CNS (n=23). Complementation crosses identified eight complementation groups and six “singleton” mutations among these 39 lines, and whole genome sequencing of most lines at ≥30X coverage identified the causative lesion(s) in and affected gene for 13 of the 14 genes. When possible, phenotypic analysis with appropriate deficiencies and gene-specific alleles was used to confirm correspondence between gene and mutant phenotype for the 13 genes. The CNS phenotype, molecular nature, and known function of these genes are summarized in Table 1 and Figure 2.

**Figure 1:**
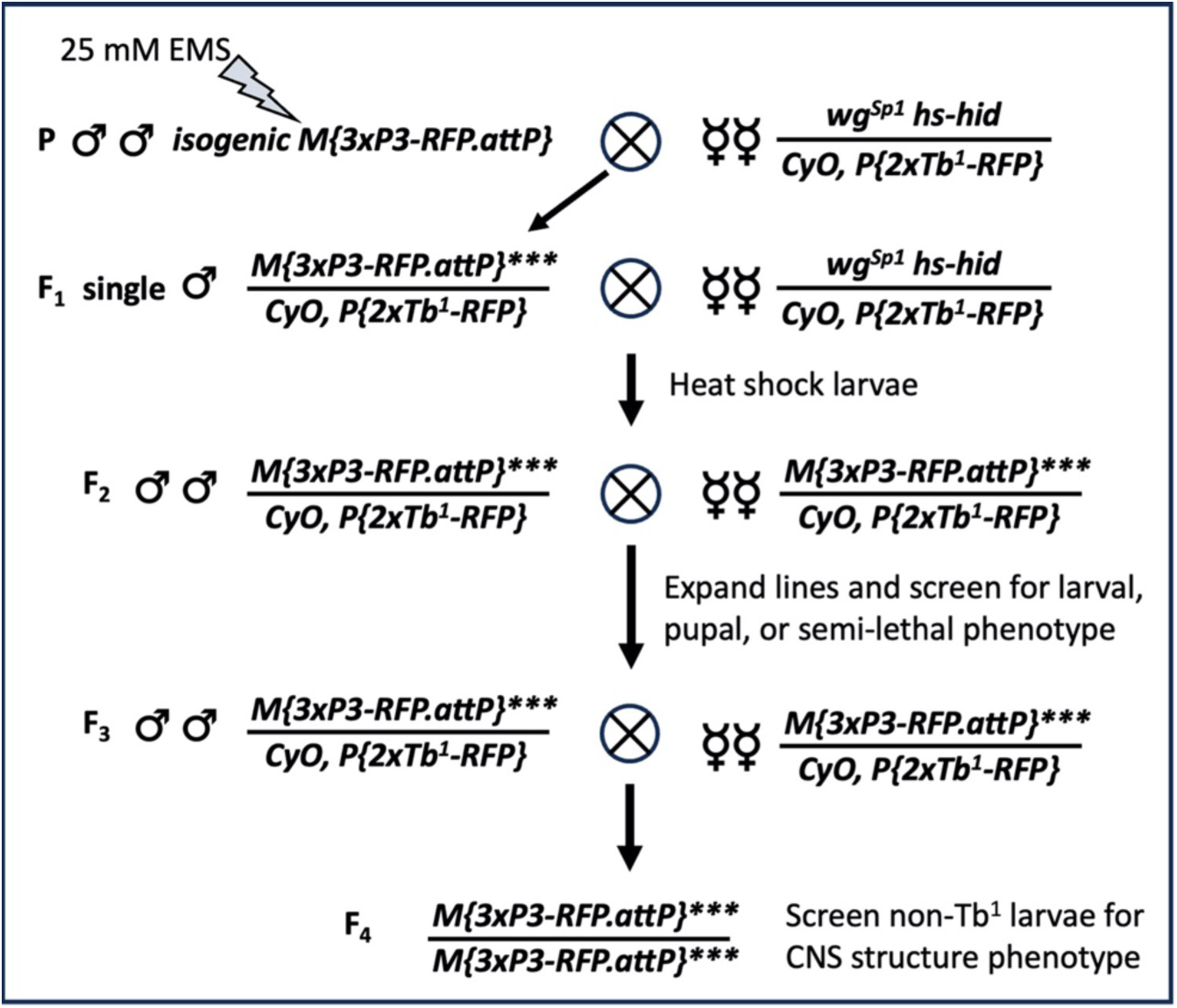
Diagram of the genetic cross scheme used to execute the forward genetic screen.

**Figure 2:**
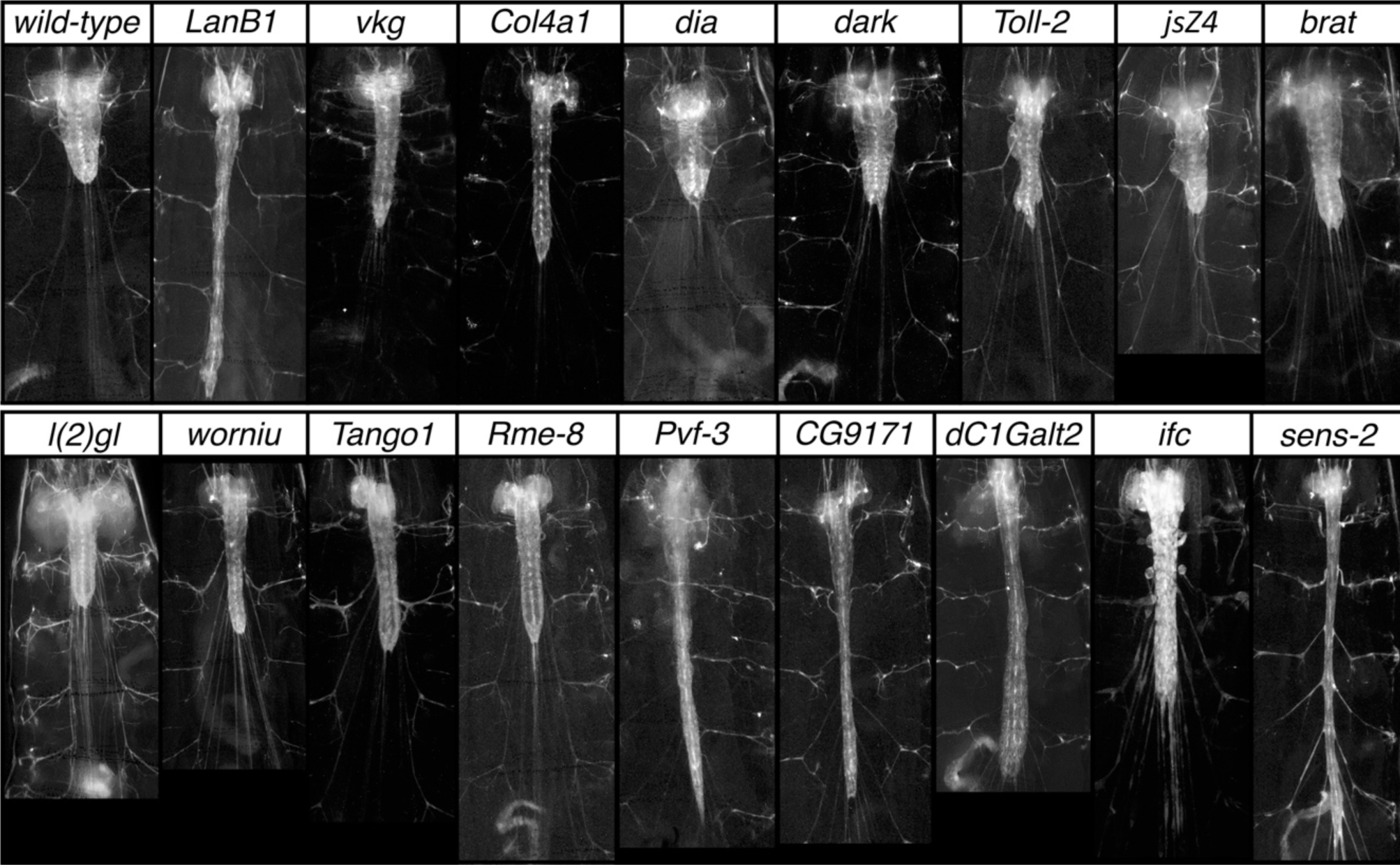
Representative examples of CNS mutant phenotypes identified in the genetic screen. Ventral views of wild-type and mutant late third instar larvae of the indicated genotype showing 3xP3 RFP staining in greyscale to highlight CNS morphology. Anterior is up; scale bar is 500μm.

**Table 1).**
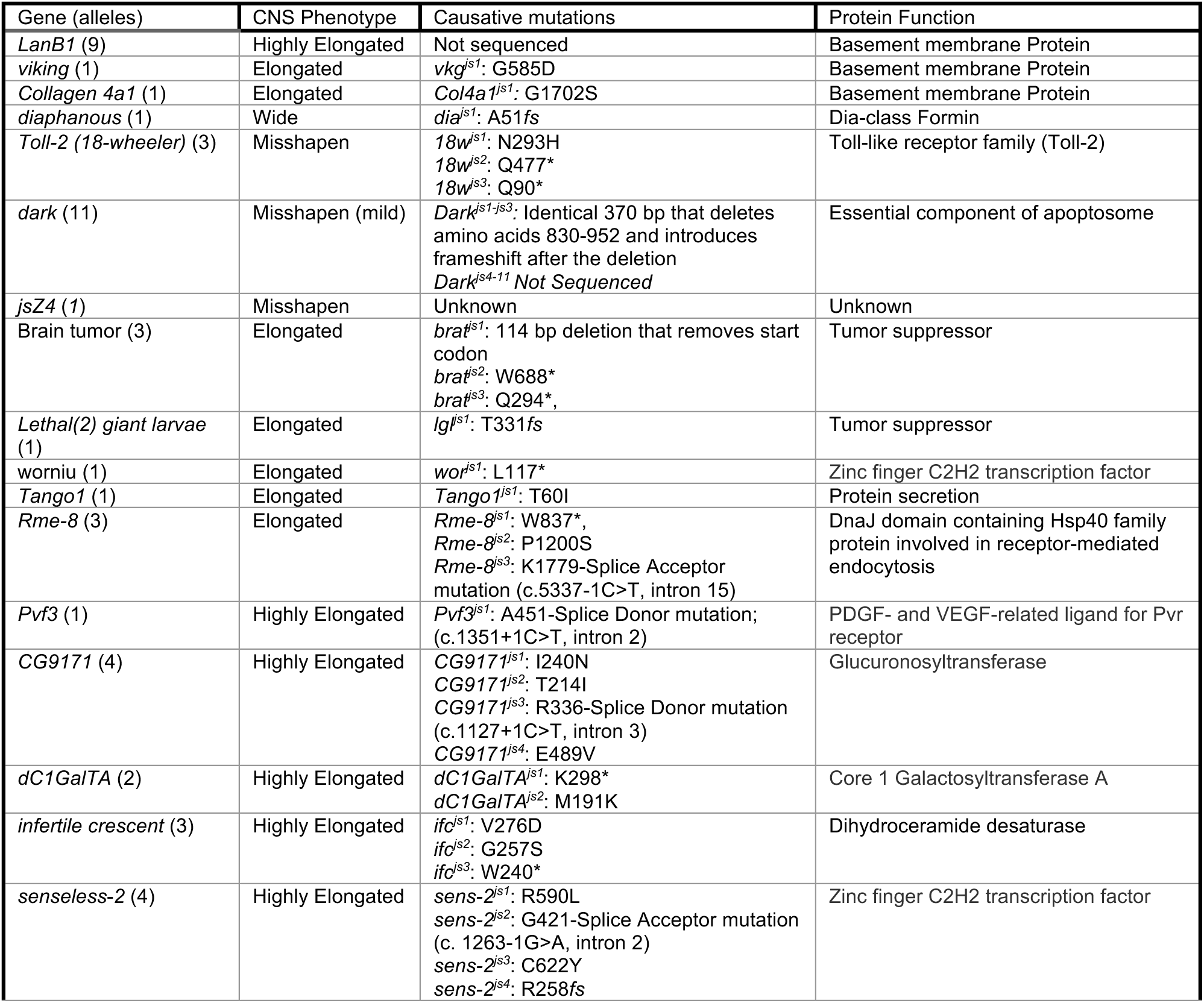
Molecular nature, phenotypic class, and causative lesions of identified genes.

Below, we outline the genes identified in the screen and their associated mutations and CNS phenotypes, starting with the gene/mutation that yielded a wider CNS and ending with those that yielded an elongated CNS. We describe in greater detail the function of the *senseless-2* gene, which encodes a Zinc finger transcription factor that may act as a genetic switch to distinguish perineurial glial found in the PNS from those in the CNS.

### Gene that yields a widened CNS phenotype when mutated

*diaphanous*: We identified one mutant allele that when homozygous yielded a wider CNS. This allele identified a frameshift mutation found immediately after the codon for amino acid 51 in the *diaphanous* coding region and resulted in an expansion of the thoracic region of the CNS that became more pronounced as larvae were about to pupariate (Table 1; Figs. 2, 3). *diaphanous* encodes the sole Dia-class formin protein in *Drosophila*; it is required for cytokinesis and induces polyploidy in follicle cells and mitotic neuroblasts (Castrillon and Wasserman 1994). Using antibodies specific to Deadpan and ELAV to label neuroblasts and neurons, respectively, we observed a massive increase in neuroblast size likely caused by defects in cytokinesis and increased ploidy of these cells (Fig. 3). Our results indicate that the expanded nature of the thoracic ventral nerve cord in *diaphanous* mutants likely results from the expanded size of neuroblasts and potentially other cell-types in the CNS.

**Figure 3:**
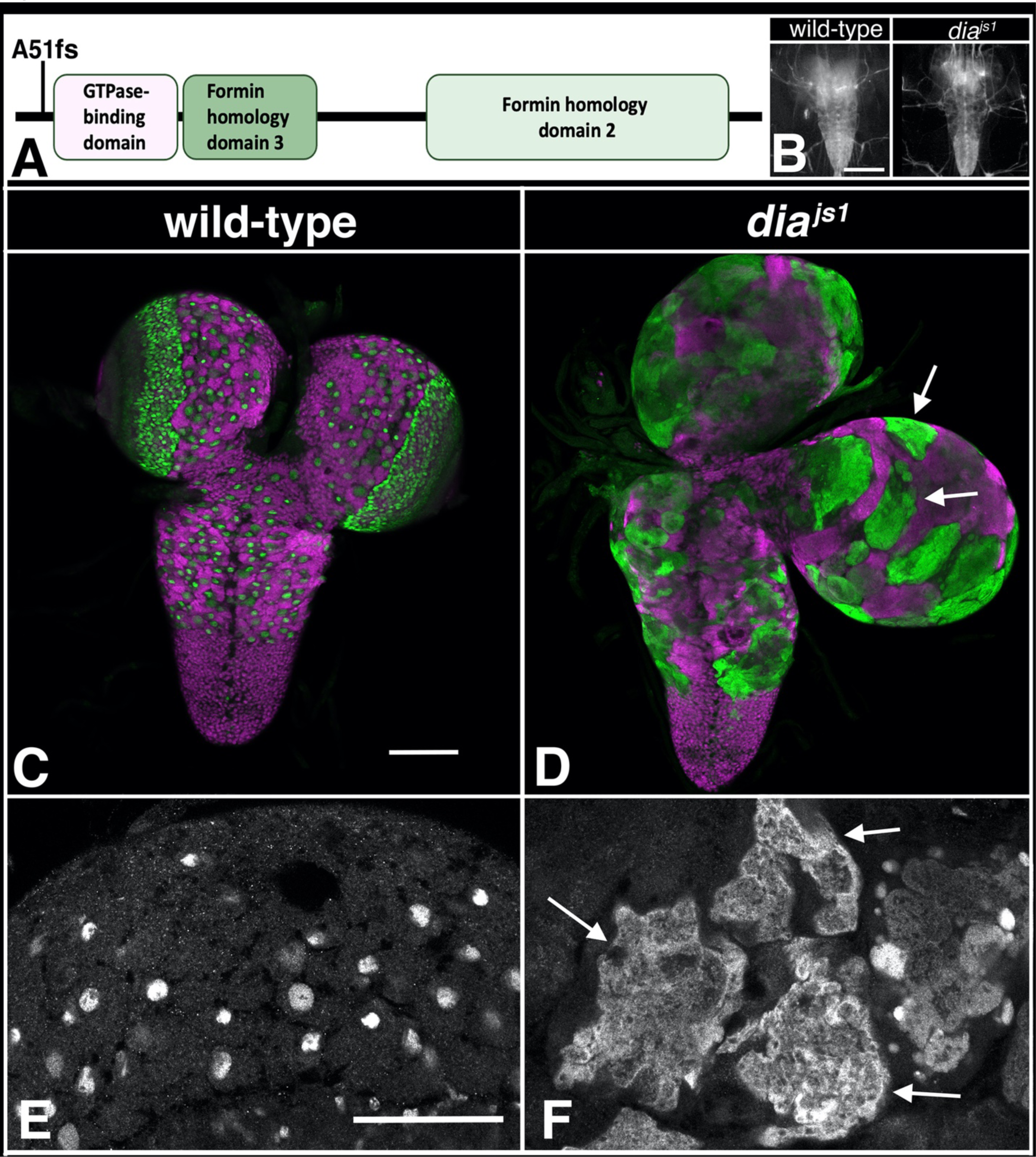
Mutations in *diaphanous* increase the width of the CNS. A) Schematic of Diaphanous protein with location and nature of the *dia^js1^* allele. B) Ventral views of wild-type or *dia^js1^* mutant late-third instar larvae with 3xP3 RFP expression (greyscale) highlighting the CNS. Anterior is up; scale bar is 250 μm. C-G) Ventral views of dissected CNS (C,D) or high magnification view of the brain (E, F) from wild-type and *dia^js1^*mutant late-third instar larvae labeled for neuroblasts with Deadpan (Dpn; green in C, D, and grey in E, F) and for neurons with ELAV (magenta in C, D). Arrows highly enlarged neuroblasts. Anterior is up; scale bar is 100 Anterior is 100 μm in panels D and E and 50 μm in F and G.

### Genes that yield a misshapen CNS phenotype when mutated

We identified mutations in three genes – *Dark, Toll-2*, and one singleton mutation – that gave rise to a misshapen CNS (Table 1). We identified 11 alleles of *Dark*. Homozygosity or trans-heterozygosity for each allele yielded a mild, partially penetrant CNS phenotype in which small hernias arise in the CNS (Fig. 2). We sequenced three of the identified alleles, each of which contained an apparently identical 370-bp deletion within the coding region (Table 1). The identity of the molecular lesions in these alleles suggest the mutations arose in mitotic spermatagonia or spermatocyte cells and are not independent of each other. *Dark* encodes a homolog of the Apaf-1/Ced-4 pro-apoptotic genes and has been shown to promote hyperplasia in the CNS (Rodriguez, Oliver et al. 1999).

#### Toll-2

Three allelic mutations identified the *Toll-2* (*18-wheeler*) gene with two of the mutations being premature stop codons and the third being a missense mutation in the leucine-rich repeat domain (Table 1; Fig. 4). Loss of *Toll-2* function reduced CNS size and resulted in a misshapen ventral nerved cord marked by irregularly shaped protrusions, hernias, and indentations (Fig. 2 and 4). The CNS phenotype of the *Toll-2* mutants grossly resembles that of *perlecan*, which encodes one of the key components of the CNS basement membrane (Yurchenco 2011), hinting that Toll-2 receptors may function in surface glia to regulate CNS morphology by interfacing with components of the basement membrane. Cell death may also contribute to the observed defects in CNS morphology, as loss of *Toll-2* function has been shown to drive cell death in neurons and neuroblasts (Li, Forero et al. 2020, Hermanstyne, Johnson et al. 2022),.

**Figure 4:**
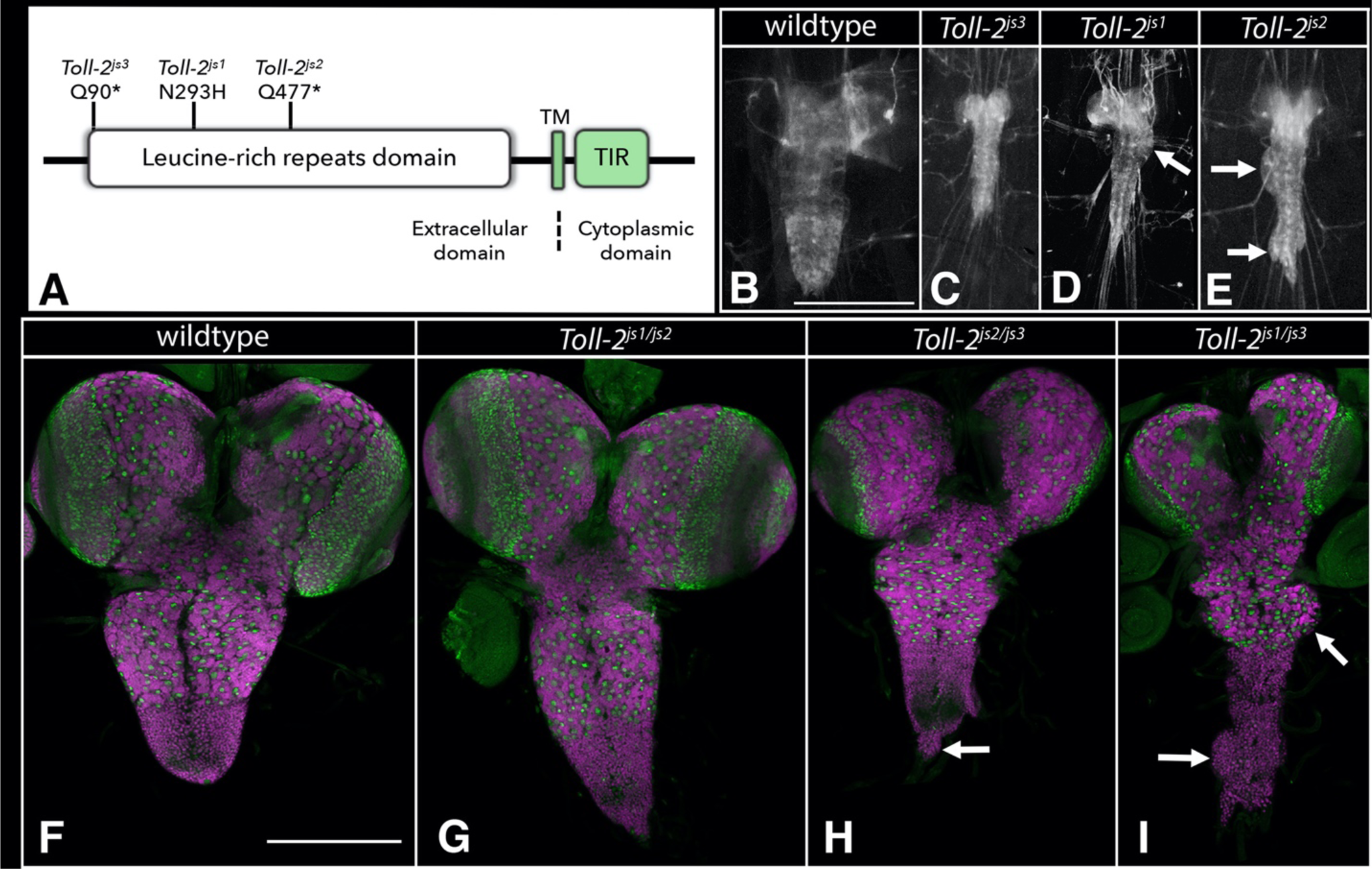
Loss of *Toll-2* function results in a misshapen CNS. A) Schematic of Toll-2 protein with location and nature of identified *Toll-2* alleles. B-E) Ventral views of wild-type or *Toll-2* mutant late-third instar larvae in which 3xP3 RFP expression (greyscale) highlights the CNS. Arrows highlight hernias, bulges, or dysmorphology in the CNS. Anterior is up; scale bar is 500μm. F-I) Ventral views of dissected CNS of wild-type (F) and *Toll-2* mutant (G-I) late-third instar larvae labeled for neuroblasts with Deadpan (Dpn; green) and neurons with ELAV (magenta). Arrows point to areas of the CNS that display dysmorphology. Anterior is up; scale bar is 200 μm.

The *jsZ4* allele represents the sole mutation that we were unable to map to a defined gene (Table 1). Its partially penetrant phenotype is highlighted by an elongated nerve cord and the presence of hernias and protrusions in it (Fig. 2). As we sequenced the genome of this allele at ≥30X coverage, our failure to map its mutant phenotype to a specific gene could arise due to the phenotype being caused by two or more mutations, a non-coding mutation, or other reasons.

### Genes that yield an elongated CNS phenotype when mutated

Our largest phenotypic class – an elongated CNS – was identified by 23 mutations and 9 genes. Two of the 9 genes are known tumor suppressors – *lethal giant larvae* (*lgl*) and *brain tumor* (*brat*). We identified a single mutation in *lgl*, a frame-shift mutation which when homozygous yields an elongated ventral nerve cord and greatly enlarged brain lobes (Table 1, Fig. 2). Complementation crosses against the *lgl^4^* allele and a deficiency that uncovers *lgl* confirmed correspondence between the gene and mutant phenotype. The elongated CNS phenotype likely arises from elevated cell proliferation since *lgl* plays a well characterized role in restricting cell proliferation (Gateff and Schneiderman 1974). Like *lgl*, loss of *brat* function is known to increase neuroblast and cell proliferation and result in tumorous growths in the brain (Gateff 1994). We identified three mutations in *brat* that yielded a moderate CNS elongation phenotype (Table 1, Fig. 2).

#### Worniu

We identified a nonsense mutation, L117*, in *worniu* that yielded a moderately elongated nerve cord (Table 1; Fig. 2). We confirmed correspondence between *worniu* and the mutant phenotype based on the new *worniu*^js1^ allele failing to complement both the *worniu^1^*and *worniu^2^* alleles for the elongated nerve cord phenotype. *worniu* encodes a C2H2-type Zinc finger transcription factor that is expressed in most neuroblasts and has been shown to regulate CNS structure (Ashraf, Ganguly et al. 2004).

#### Tango1

We identified a missense mutation, T60I, in the *Tango1* coding region that when homozygous resulted in a moderately elongated CNS (Table 1; Fig. 2). We confirmed correspondence between this mutation and the observed CNS elongation phenotype by complementation against two overlapping deficiencies that each uncover *Tango1*. *Tango1* functions at ER exit sites to promote protein secretion and is required to promote the specialized secretion of the large Collagen and Laminin proteins (Saito, Chen et al. 2009, Malhotra and Erlmann 2011, Wilson, Phamluong et al. 2011, Petley-Ragan, Ardiel et al. 2016). Thus, the CNS elongation phenotype of *Tango1* likely derives from defective secretion and deposition of basement membrane proteins, as loss of function in the basement membrane proteins, such as *LanB1*, *vkg*, and *Col4a1* lead to an elongated CNS (Pastor-Pareja and Xu 2011, Kim, Jeibmann et al. 2014, Skeath, Wilson et al. 2017, Dai, Estrada et al. 2018).

#### Rme-8

Three mutations in the *Rme-8* gene also gave rise to a moderately elongated CNS phenotype when homozygous or trans-heterozygous to each other (Table 1; Fig. 1, 5). Rme-8 encodes a DnaJ domain-containing protein of the Hsp40 chaperone family that also contains four IWN repeat (Zhang, Grant and Hirsh 2001, Norris, McManus et al. 2022); Rme-8 promotes endocytic recycling of transmembrane proteins, such as Notch (Gomez-Lamarca, Snowdon et al. 2015). Using an *Rme-8-T2A-GAL4* CRIMIC insert in the fourth intron of *Rme-8* to drive a UAS-linked nuclear-RFP transgene (Lee, Zirin et al. 2018, He, Binari et al. 2019), we found that Rme-8 is broadly expressed in the CNS in neurons and glia, especially surface glia (data not shown).

The remaining five genes (*Pvf3, CG9171, C1GalTA, ifc,* and *sens-2*) all exhibited severe CNS elongation phenotypes upon loss of their function.

#### Pvf3

We identified a single mutation in *Pvf3* – a splice site donor mutation just prior to the region that encodes its PDGF domain (Table 1; Fig. 2, 6). Larvae homozygous for this mutation yield a highly elongated CNS, which is recapitulated when this allele is placed in trans to the *Df(2L)τ.Pvf2-3* deficiency or two P element inserts in *Pvf3* (see Methods; (Parsons and Foley 2013)), confirming correspondence between *Pvf3* and the elongated CNS phenotype. *Pvf3* acts together with *Pvf2* to promote hemocyte survival and migration through activation of the Pvr receptor (Parsons and Foley 2013). Loss of Pvr function or blockade of its neural activity has been shown to inhibit nerve cord condensation and promote nerve cord elongation (Olofsson and Page 2005). Our observation that loss of *Pvf3* function alone yields a highly elongated CNS phenotype suggests that *Pvf3* plays a non-redundant role relative to *Pvf2* to promote hemocyte survival, migration, and/or function.

#### CG9171

Four mutant alleles identified *CG9171*, which encodes a predicted glucuronosyltransferase that promotes O-linked protein mannosylation. All four mutations reside in its extracellular glycosyl-transferase domain, likely disrupting enzymatic function (Fig. 7). Homozygous or trans-heterozygous mutants of all four mutations yield a highly elongated CNS phenotype, but neuroblast formation and neuronal patterning appear grossly normal (Fig. 7). The most closely related mammalian homologs of *CG9171* – *B4GAT1* and *LARGE1/2* – act in tandem to drive the O-mannosylation of Dystroglycan (Praissman, Live et al. 2014). B4GAT1 adds a single glucuronic acid residue onto a Xylose acceptor on Dystroglycan. The related glycosyltransferase LARGE recognizes this epitope and extends it by adding many copies of a repeating disaccharide to form matriglycan. Matriglycan links Dystroglycan to the basement membrane by binding to Laminin and other proteins and plays a key role in disease, as mutations in B4GAT1 in humans cause dystroglycanopathies likely due to the loss of matriglycan on Dystroglycan (Yoshida-Moriguchi, Yu et al. 2010, Praissman, Live et al. 2014, Bigotti and Brancaccio 2021). Our work suggests that defects in the O-mannosylation of Dystroglycan and/or other cell surface proteins cause the observed CNS elongation phenotype.

**Figure 5:**
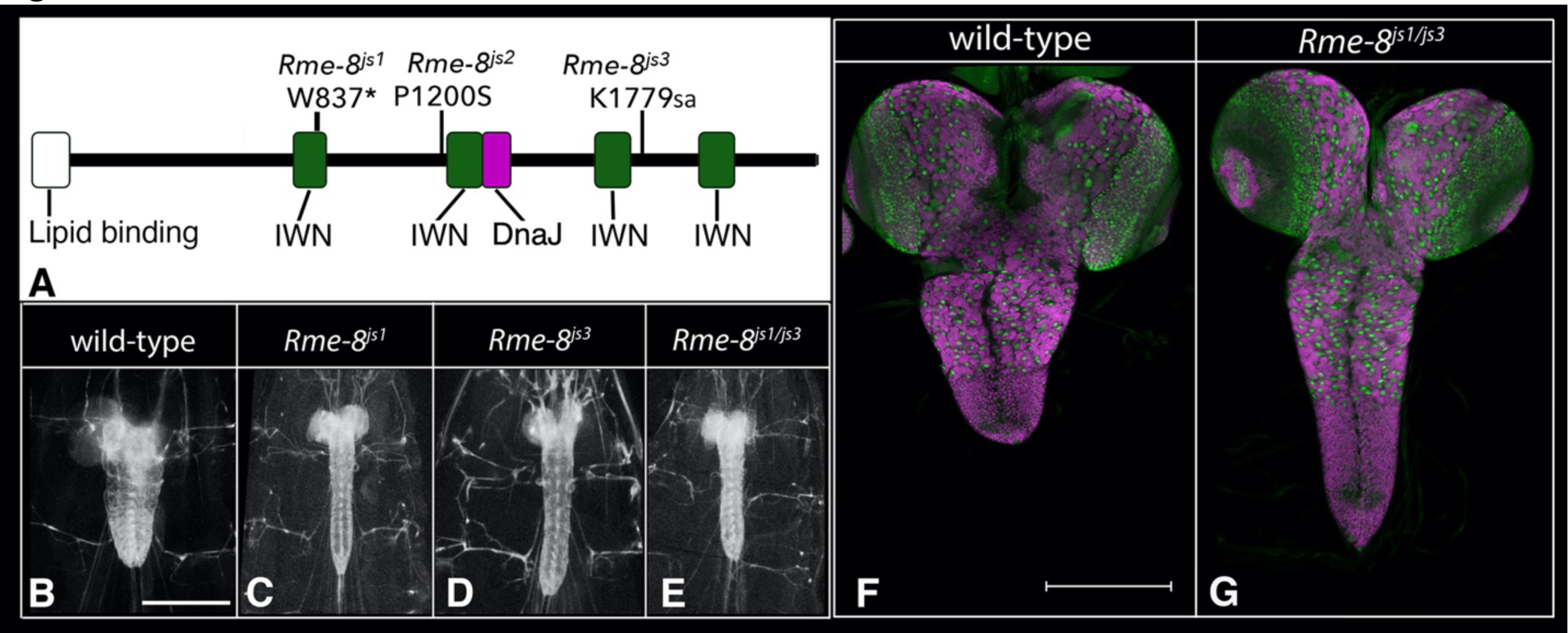
Loss of *Rme-8* function promotes CNS elongation. A) Schematic of Rme-8 protein showing the location of the lipid binding, IWN repeat, and DnaJ domains as well as the location and nature of the three identified *Rme-8* alleles. B-E) Ventral views of wild-type or *Rme-8* mutant late-third instar larvae in which 3xP3 RFP expression (greyscale) highlights the CNS. Anterior is up; scale bar is 500μm. F-G) Ventral views of dissected CNS of wild-type (F) and *Rme-8* mutant (G) late-third instar larvae labeled for neuroblasts with Dpn (green) and neurons with ELAV (magenta). Anterior is up; scale bar is 200 μm.

**Figure 6:**
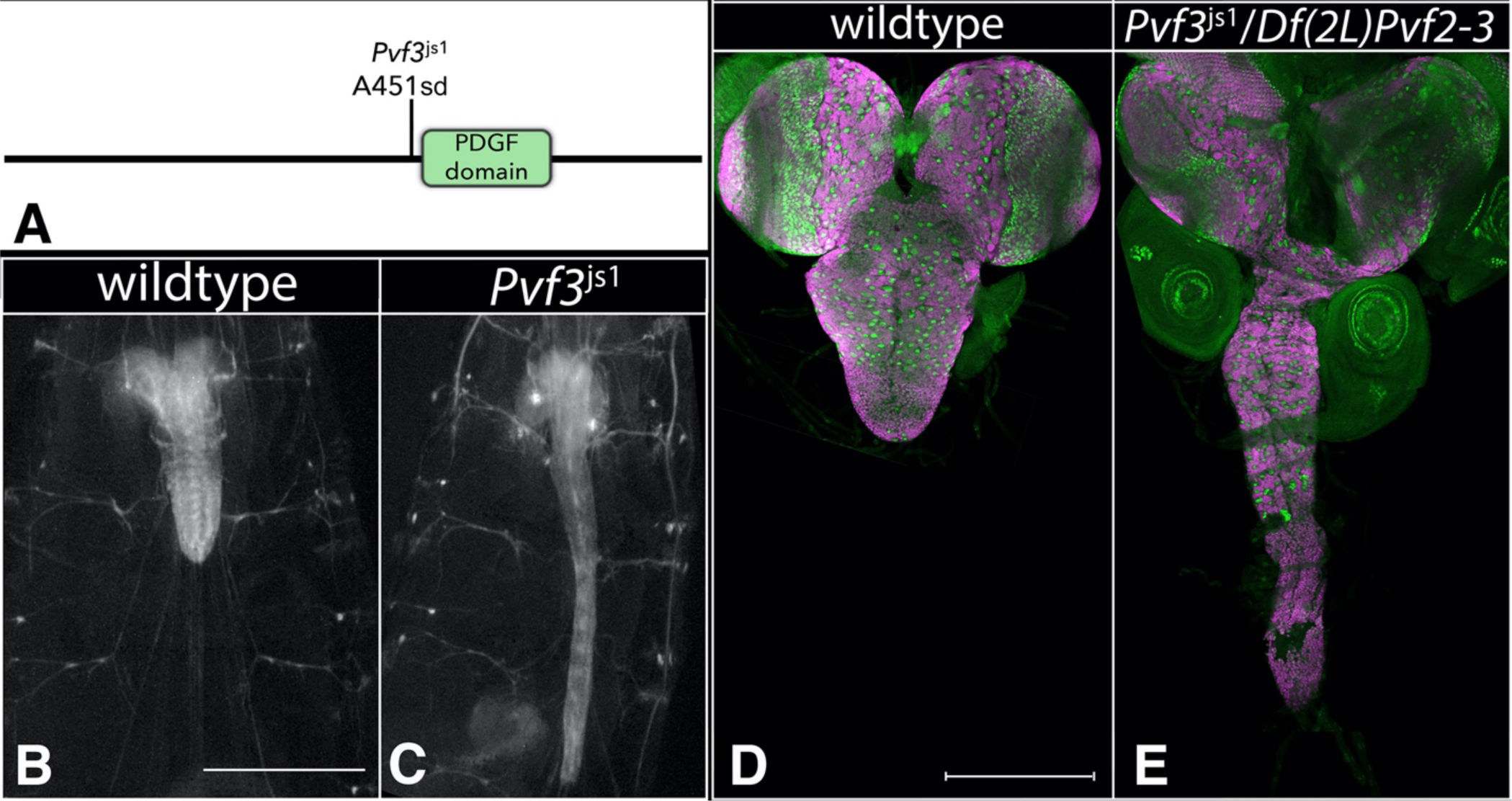
Loss of *Pvf-3* function greatly increases the length of the ventral nerve cord. A) Schematic of Pvf-3 protein with location and nature of the *Pvf-3^js1^* allele. B-E) Ventral views of wild-type or *Pvf-3^js1^* mutant late-third instar larvae in which 3xP3 RFP expression (greyscale) highlights the CNS. Anterior is up; scale bar is 500μm. D, E) Ventral views of dissected CNS of wild-type (F) and *Pvf-3^js1^* mutant (G-I) late-third instar larvae labeled for neuroblasts with Deadpan (Dpn; green) and for neurons with ELAV (magenta). Anterior is up; scale bar is 200 μm.

**Figure 7:**
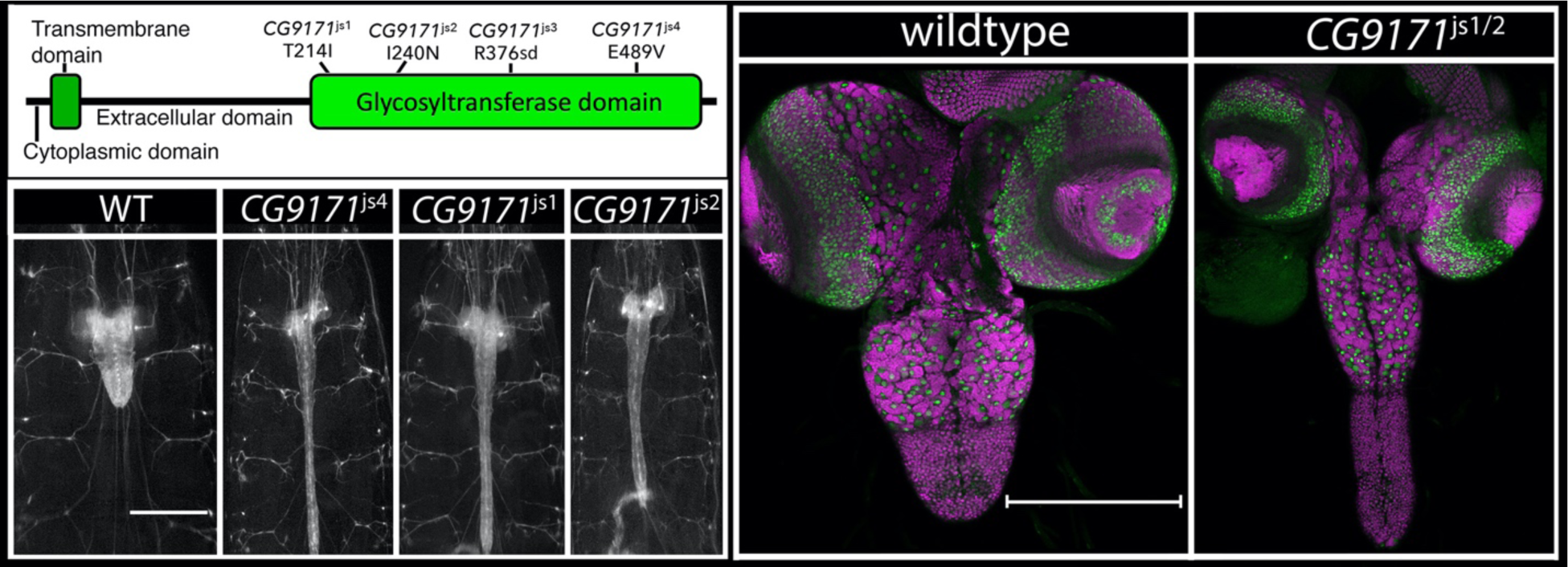
Loss of *CG9171* drives CNS elongation. A) Schematic of CG9171 protein with location and nature of the four identified *CG9171* alleles. B-F) Ventral views of wild-type or *CG9171* mutant late-third instar larvae in which 3xP3 RFP expression (greyscale) highlights the CNS. Anterior is up; scale bar is 500μm. G-H) Ventral views of dissected CNS of wild-type (G) and *CG9171* mutant (H) late-third instar larvae labeled for neuroblasts with Dpn (green) and neurons with ELAV (magenta). Anterior is up; scale bar is 200 μm.

#### C1GalTA

Like *CG9171*, the two alleles in *Core 1 Galactosyltransferase A* (*C1GalTA; CG9520*) yielded a highly elongated CNS phenotype (Table 1; Fig. 2). Unlike the CG9171 phenotype in which the mutant ventral nerve cord is supple and flexible, *C1GalTA* mutant larvae exhibited a highly elongated, very brittle nerve cord, flat and misshapen brain lobes, reduced fat body tissue integrity, and direct adherence of portions of the fat body to the CNS (Fig. 8; data not shown). These phenotypes are most obvious in larvae homozygous for the phenotypically stronger *C1GalTA^js1^* allele. C1GalTA promotes protein glycosylation by adding galactose in a β1,3 linkage to N-acetylgalactosamine (GalNAc) on proteins. In flies, a prior study showed that *dC1GalTA* is required for glycosylation of Laminin and the presence of T antigen (Gal β1,3 GalNAc) on hemocytes, and that loss of *C1GalTA* function drives CNS elongation (Lin, Reddy and Irvine 2008).

**Figure 8:**
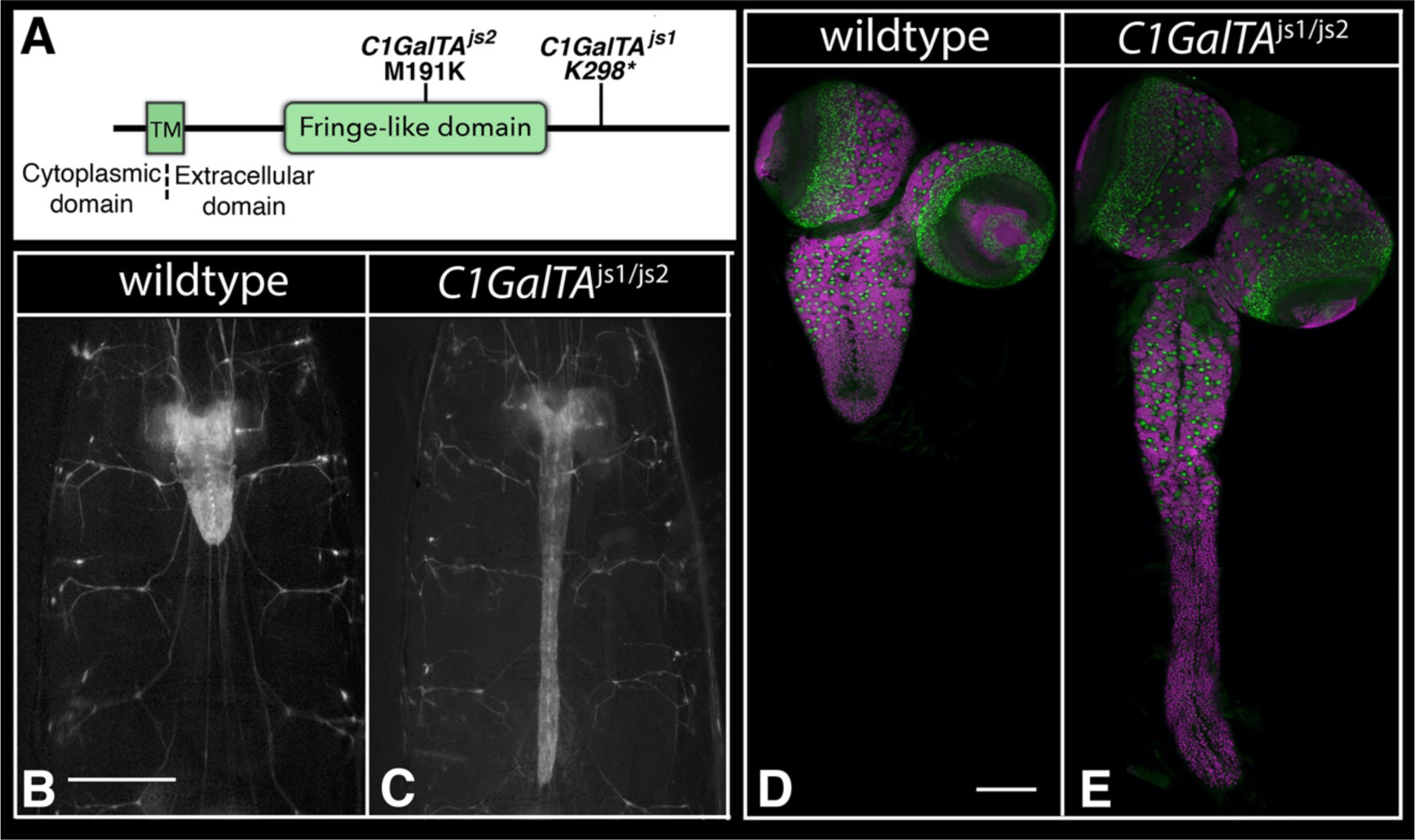
Loss of *dC1GalTA* function promotes CNS elongation. A) Schematic of C1GalTA protein with location of its Fringe-like glycosyltransferase domain and the nature of the two identified alleles. B-C) Ventral views of wild-type or *C1GalTA^js1/js2^* mutant late-third instar larvae in which 3xP3 RFP expression (greyscale) highlights the CNS. Anterior is up; scale bar is 500μm. D, E) Ventral views of dissected CNS of wild-type (F) and *C1GalTA^js1/js2^* mutant (G-I) late-third instar larvae labeled for neuroblasts with Deadpan (Dpn; green) and for neurons with ELAV (magenta). Anterior is up; scale bar is 100 μm.

The *C1GalTA* mutant phenotype resembles that of *papilin* (*ppn*), a third chromosomal gene that we were working on in parallel to the genetic screen due to its sequence similarity to AdamTS genes (Campbell, Fessler et al. 1987, Kramerova, Kawaguchi et al. 2000, Keeley, Hastie et al. 2020). Ppn is a key component of the basement membrane, where it has been shown to facilitate Collagen IV removal to promote basement membrane remodeling (Keeley et al., 2020). Like the *C1GalTA* mutant phenotype, reduction of *ppn* function yields a highly elongated and brittle nerve cord, flat and misshapen brain lobes, reduced fat body integrity, and portions of the fat body adhering to the CNS (Fig. 9, data not shown). A *ppn*-T2A-GAL4 reporter and Ppn-specific antibodies showed that *ppn* is transcribed in hemocytes, the cells that assemble and disassemble basement membranes, but not in most other tissues (Fig. 9; data not shown), including the fat body – the major producer of basement membrane proteins in larvae (Pastor-Pareja and Xu 2011). Ppn protein, however, is found on the basement membrane of most tissues (not shown). We hypothesize that hemocytes deposit Ppn on most tissues during development.

**Figure 9:**
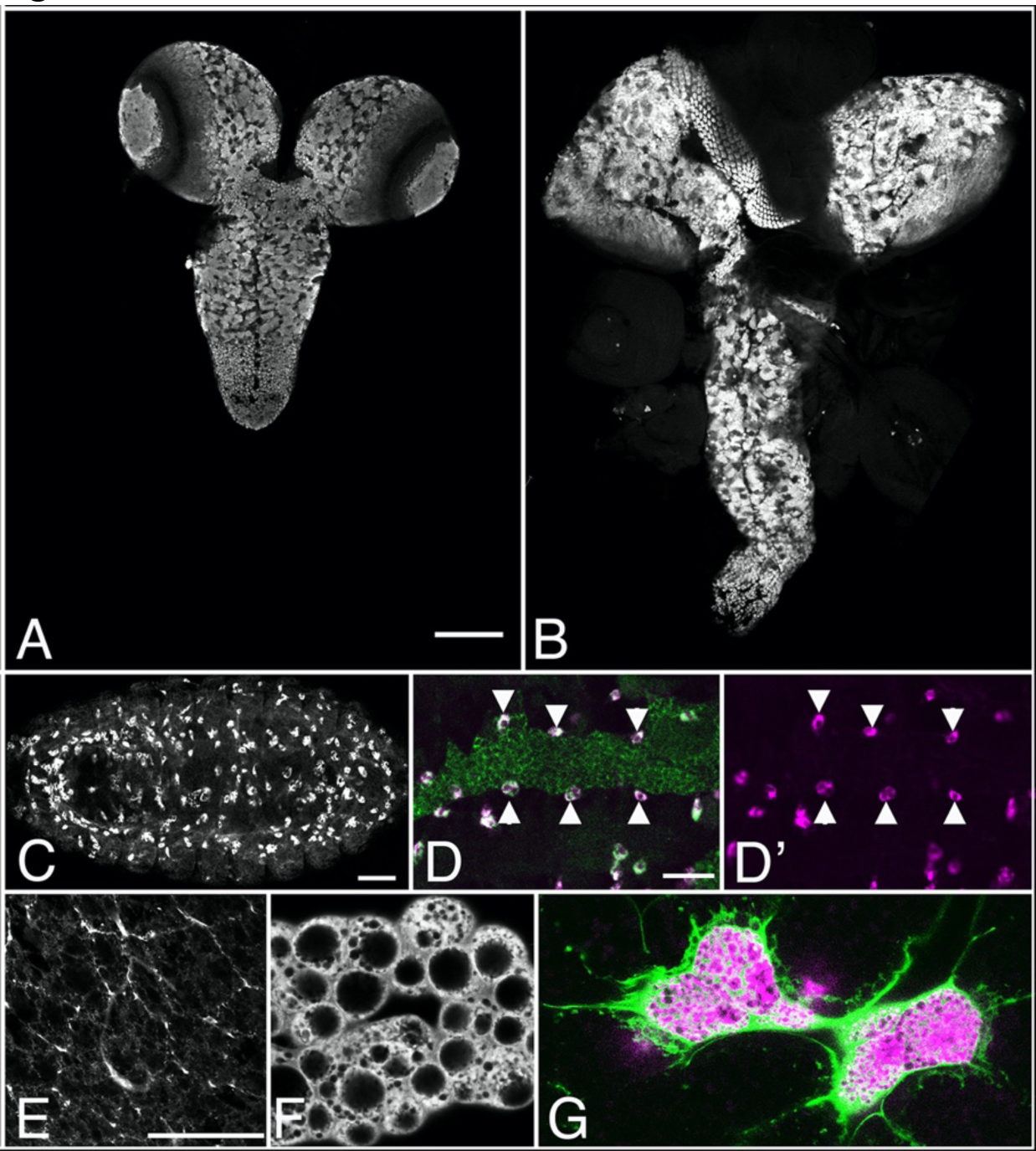
*papilin* regulates CNS structure and is sufficient to drive accumulation of Vkg-GFP in the fat body. A-B) Ventral view of CNS of wild-type (A) and *ppn^MI03189^* mutant late third instar larvae labeled for ELAV (grey scale). Anterior is up; scale bar is 100 μm. (C-D) Stage 15-16 embryos labeled for Ppn (grey, C; red, D-D’) and Collagen 4a1 (green, D). Anterior is to the left; scale bars are 50 μm. Arrowheads in D-D’ point to Ppn^+^ hemocytes immediately adjacent to the fat body. E-G) vkg-GFP localization (grey in E, F; green in G) in wild-type fat body (E), fat body in which Ppn[RG] is expressed in all fat body cells (F), and fat body in which small cell clones, marked in magenta, ectopically express Papilin protein (G). Scale bar is 50 μm.

Most basement membrane components, like Col4a1, Vkg, Nidogen, and Perlecan, are produced by fat body cells (Pastor-Pareja and Xu 2011), but Ppn is not expressed in the fat body, suggesting there may be a physiological reason for exclusion of Ppn transcription in the fat body. To test this model, we misexpressed *ppn* in the fat body and asked if it altered basement membrane protein localization by visualizing the localization of a GFP protein trap in Viking (Morin, Daneman et al. 2001). Upon forced *ppn* expression in the fat body, we observed a massive, cell autonomous retention of Viking-GFP in fat body cells (Fig. 9), demonstrating incompatibility of fat body *ppn* expression with appropriate Viking secretion and suggesting that Ppn physically associates with Viking in vivo to help mediate the removal of the basement membrane components. The similarity of the *ppn* and *C1GalTA* mutant phenotypes suggests that C1GalTA acts with Ppn to promote Viking/Collagen IV removal from the basement membrane and that failure to do so results in a brittle, elongated CNS. Future work that determines how these genes functionally interface with each other should help clarify the molecular basis of basement membrane remodeling and the control of tissue structure.

#### infertile crescent

Three alleles identified *infertile crescent (ifc)*, which encodes the sole *Drosophila* dihydroceramide desaturase that converts dihydroceramide to ceramide in the de novo ceramide biosynthesis pathway (Jung, Liu et al. 2017) (Hannun and Obeid 2018). Loss of *ifc* results in an elongated CNS marked by bulges in peripheral nerves (Fig. 2); we characterized the function of *ifc* in detail in a separate study and found it primarily functions in glia to promote glial homeostasis and to protect against neuronal death and neuronal degeneration (Zhu, Cho et al. 2024).

### Senseless-2 acts in perineurial glia to regulate CNS structure

Four mutant alleles identified the *senseless-2* (*sens-2*) gene, which encodes a member of the Zinc finger C2H2 superfamily of proteins and possesses six C2H2 Zinc finger domains (Fig. 2, 10A). Three of the alleles – *sens-2^js2^, sens-2^js3^,* and *sens-2^js4^* – introduce, respectively, an early splice site mutation, a missense mutation in a conserved cysteine in the Zinc finger domain, and an early frame-shift mutation (Table 1). These alleles yield an essentially identical highly elongated CNS phenotype (Fig. 10B, 10C) and are then likely null or strong hypomorphic alleles. The fourth allele, *sens-2^js1^*, yields a modest CNS elongation phenotype; larvae trans-heterozygous for *sens-2^js1^*and *sens-2^js4^* exhibit an intermediate CNS elongation phenotype between the two alleles, identifying *sens-2^js1^*, which contains a missense mutation in the zinc finger domain (Table 1), as a weak hypomorphic allele of *sens-2*.

**Figure 10:**
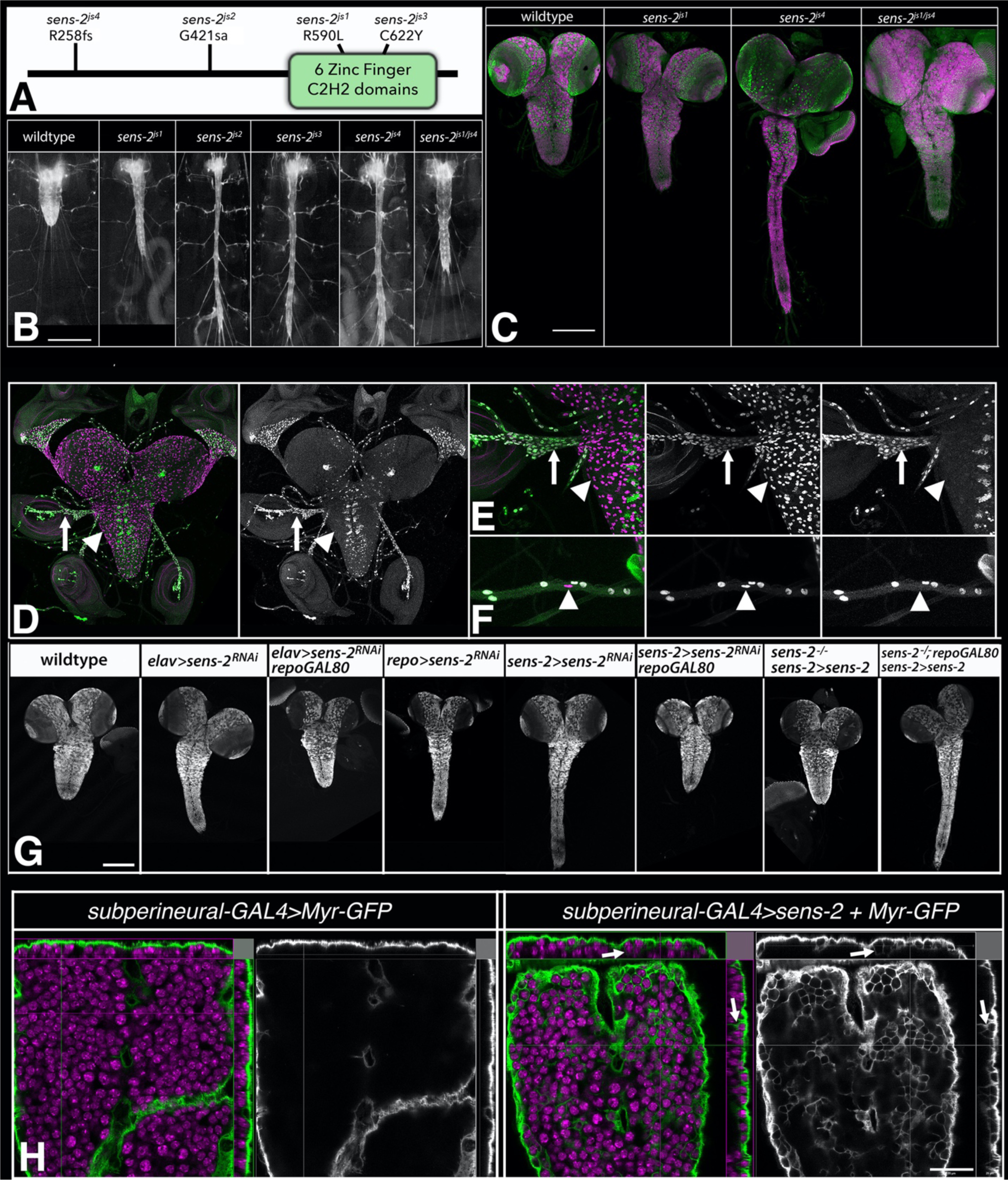
*sens-2* acts in peripheral glia to regulate CNS structure. A) Schematic of Sens-2 protein with location and nature of the four identified *sens-2* mutations. B) Ventral views of late-third instar larvae of indicated genotype showing 3xP3 RFP expression (greyscale) in the CNS and nerves. Anterior is up; scale bar is 500μm. C) Ventral views of photomontages of late third instar larvae of the indicated genotypes labeled for neuroblasts with Deadpan (Dpn; green) and for neurons with ELAV (magenta). Anterior is up; scale bar is 200 μm. D-F) Ventral views of low (D) and high magnification (E,F) images of the CNS (D,E) and peripheral nerves (F) of late third instar larvae labeled for *sens-2-GFP* (green and greyscale) to mark *sens-2* expressing cells and REPO (magenta) to mark glia. Arrows in C,D identify peripheral glia that express *sens-2-GFP*; arrowheads identify central glia that lack GFP expression. Arrowhead in E marks *sens-2-GFP*-negative wrapping glia in peripheral nerve. G) Ventral views of the dissected CNS from late third instar larvae of indicated genotype labeled for ELAV (greyscale) to mark neurons and highlight the structure of the CNS. Anterior is up; scale bar is 200 μm. H) Ventral view and Y-Z and X-Z projection views of late third instar larvae of indicated genotype labeled for Myr-GFP (green/greyscale) and ELAV to label neurons (magenta). Arrows point to Myr-GFP+ membrane invaginations that enwrap neurons. Anterior is up; scale bar is 20 μm.

The *sens-2* CNS elongation phenotype manifests in first instar larvae and is maintained throughout larval life and into pupal stage at which point *sens-2* homozygous flies die. In live third instar larvae, the CNS appears to be under tension from both laterally and posteriorly projecting peripheral nerves, with the posterior nerves being shorter in length in *sens-2* mutant larvae relative to wild-type (Fig. 2, 10A). The overall shape of the CNS is dramatically altered, but the overall pattern of neuroblasts and neurons appeared grossly wild-type with no significant change in neuroblast number (Fig. 10A).

To clarify the cellular basis for the *sens-2* CNS elongation phenotype, we tracked *sens-2* expression in larvae by generating a Sens-2 specific antibody and a *sens-2-T2A-GAL4* CRIMIC line (“*sens-2-GAL4*”) and leveraging a *sens-2-GFP* transgene surrounded by its endogenous genomic locus ((Kudron, Victorsen et al. 2018); see Methods). These tools revealed that *sens-2* is expressed in subsets of glia and neurons in the larval nervous system (Figs. 10C-F, S1-3) and in a region-specific manner in the gut (not shown). We focused on *sens-2* expression in the nervous system, as its function in glia, neurons, or both likely contributes to the observed CNS phenotype. Double-labeling experiments with REPO to mark all glia and ELAV to label all neurons revealed that *sens-2* is expressed in a small number of CNS neurons and most glia in peripheral nerves, but that *sens-2* is not detected in any glia within the CNS itself (Figs. 10D-E, S2, S3). More detailed analysis of *sens-2* expression within peripheral nerves of late third instar larvae suggested that *sens-2* is expressed in all perineurial glia and weakly in some subperineural glia, but that its expression appears to be excluded from wrapping glia and most subperineurial glia (Figs. 10F, S2, S3). We conclude that *sens-2* expression demarcates PNS perineurial glia (*sens-2*-positive) from CNS perineurial glia (*sens-2*-negative), identifying *sens-2* as one of the first markers to distinguish a glial subtype along the CNS-PNS axis.

To determine if *sens-2* function is required in glia, neurons or both to regulate CNS structure, we used the GAL4/UAS system together with either a *UAS-sens-2-RNAi* transgene or a *UAS-sens-2* transgene (Fig. 10G). Depletion of *sens-2* function specifically in neurons via RNAi using *elav-GAL4* together with *repo*-*GAL80* had no effect on CNS structure or morphology (n=11/11; Fig. 10G), but depleting *sens-2* function specifically in glia using *repo-GAL4* recapitulated the elongated nerve cord phenotype observed in *sens-2* mutant larvae (n=11/11; Fig. 10G). We note that using *elav-GAL4* in the absence of *repo-GAL80* also resulted in elongated nerve cords (n=11/12) due to the ability of *elav-GAL4* to drive significant gene expression in glia (Berger, Renner et al. 2007). We observed similar results with the *sens-2-GAL4* line: RNAi-mediated depletion of *sens-2* in all *sens-2* expressing cells elicited an elongated nerve cord (n=12/12), but when *repo-GAL80* was used to block *sens-2* RNAi in glia, the phenotype disappeared, and nerve cords appeared wild-type (n=11/11). Gene rescue experiments corroborated the above results: Expression of a wild-type *sens-2* transgene under control of the *sens-2-GAL4* line in otherwise *sens-2* mutant larvae fully rescued the *sens-2* CNS phenotype (n=10/10), but when *repo-GAL80* was used to block transgene expression specifically in glia, the *sens-2* elongated CNS phenotype reappeared (n=10/10). We conclude that *sens-2* function is required solely in PNS perineurial and perhaps subperineural glia to control CNS structure, and that loss of *sens-2* function in glial cells alone is sufficient to yield the observed elongated nerve cord phenotype.

To assess the function of *sens-2* in different glial subtypes, we leveraged the GAL4/UAS system and GAL4 lines specific for each glial subtype to drive *sens-2* expression and that of *Myr-GFP*, which outlines cell morphology, in each glial subtype. We observed no gross change in cell morphology upon *sens-2* misexpression in astrocyte-like, cortex, and ensheathing glia. Forced expression of *sens-2* by two different perineurial-specific GAL4 lines, which also drive strong gene expression in the gut, resulted in early larval lethality inhibiting our ability to assess the impact of *sens-2* misexpression in this glial subtype. Misexpression of *sens-2* in subperineural glia, however, drove a clear change in cell morphology. Subperineural glia normally form a thin, flat cell layer that fully encircles the circumference of the CNS and peripheral nerves and resides immediately interior to the perineurial glial cell layer (Yildirim, Petri et al. 2019). Upon *sens-2* misexpression in subperineural glia, these glial cells still fully encircle the CNS, but they now also extend cell membranes into the interior of the CNS to fully or partially enwrap individual neurons (arrows, Fig. 10H). *sens-2* misexpression in subperineural glia then modifies the behavior of this glial cell type, bestowing on it the ability to enwrap neuronal cell bodies in addition to the entire CNS, suggesting that *sens-2* alters the membrane properties of surface glia to facilitate their ability to wrap peripheral nerves.

Our work on *sens-2* suggests that it acts as a genetic switch to distinguish the functional properties of perineurial glia found In the PNS from those found in the CNS. A key difference between these two tissues is their diameter – peripheral nerves are tiny in diameter relative to the much larger CNS. In this context, the ability of forced *sens-2* expression to alter the behavior of subperineural glia so that they inappropriately enwrap adjacent neuronal cell bodies highlights a profound impact of *sens-2* expression on key functional properties of surface glia – their ability to enwrap (or not enwrap) small diameter structures, such as cells or nerves. Our work did not clarify the molecular basis through which *sens-2* dictates such functional properties. In the future it will be important to identify the transcriptional targets of *sens-2* to clarify the exact mechanism through which it governs the development and differentiation of perineurial glia in the PNS.

Summary: Prior work from many labs has highlighted the importance of interactions between the basement membrane and glial cells in dictating CNS structure (Kim, Jeibmann et al. 2014, Meyer, Schmidt and Klambt 2014, Skeath, Wilson et al. 2017). Initial results from our screen reinforce these findings. *Tango1, C1GalTA, ppn,* and *pvf3* all appear to act on basement membrane proteins, form components of the basement membrane, or regulate the survival or migration of hemocytes, the bricklayers and tuckpointers of the basement membrane. Continued work on these genes, especially *C1GalTA* and *ppn*, which likely act in the same pathway, holds the promise of clarifying our understanding of basement membrane function and remodeling on tissue structure. Conversely, *sens-2* and *ifc* act in glial cells to regulate CNS structure (this paper, Zhu et al. 2024). We expect *Rme-8, CG9171*, and *Toll-2* also act in glia to regulate CNS shape, with future work needed to confirm this prediction and to clarify if and how such factors interface with the basement membrane to govern CNS morphology.

## Materials and methods

### Genetic screen

To identify mutations that disrupt larval CNS morphology, we performed a standard, autosomal recessive forward genetic screen of the second chromosome using a chromosome isogenic for the *M[3xP3-RFP.attP]* ϕC31 “landing pad” transgene inserted in cytological location 51D. The *3xP3-RFP* construct in this transgene expresses RFP in most glia and allows rapid visualization of CNS morphology in live larvae (Zhu, Cho et al. 2024). As outlined in Figure 1, we mutagenized isogenic *M[3xP3-RFP.attP]* males with 25mM EMS for 8 hours, allowed them to recover from EMS treatment overnight, and then mated them *en masse* to *CyO P[Tb^1^-RFP]/wg^Sp1^ P[Hs-hid]* virgin females, discarding mutagenized males on day 5. The CyO *P[Tb^1^-RFP]* chromosome carries the larval marker *Tubby* (*Tb*), facilitating the identification of larvae that carry or do not carry this balancer based on larval body shape. The *wg^Sp1^ P[Hs-hid]* chromosome carries the *hid* transgene under the control of the *hsp70* promoter, which induces activation of the pro-apoptotic *hid* gene upon heatshock, killing all flies that carry this transgene. 17,790 F1 males of the *M[3xP3-RFP.attP]***/*CyO *P[Tb^1^-RFP]* genotype, where “***” denotes the mutagenized chromosome, were individually crossed to 3-5 *CyO P[Tb^1^-RFP]/wg^Sp1^ P[Hs-hid]* virgin females and allowed to mate for 7 days before adults were discarded. On days 8 and 9, each vial was heat-shocked for 30-35 minutes at 37°C to ensure that only F2 progeny of the following genotype survived: *M[3xP3-RFP.attP]***/*CyO *P[Tb^1^-RFP].* 12,211 of the 17,790 single male crosses were successful; for each of these crosses, F2 progeny were sib-mated to expand the stock and resulting F3 progeny were screened for larval-, pupal-, or semi-lethal phenotypes. 3180 lines displayed such a phenotype. For each of these lines, F4 larvae homozygous for the mutagenized chromosome, easily identified by their lack of the *Tb* marker, were visually screened for disrupted CNS morphology, e.g. an elongated CNS, under a fluorescent dissecting microscope. 50 lines were identified that harbored second chromosomal mutations that when homozygous disrupted the morphology of the larval CNS.

### Complementation Analysis

The 50 mutations grouped into three phenotypic classes: those with an elongated CNS (n=34), a herniated or mis-shapen CNS (n=15), or a wider CNS (n=1). Exhaustive complementation crosses among lines within each group identified nine complementation groups and nine “singleton” mutations. Crosses of each of these lines to second chromosomal mutations known to disrupt CNS morphology, such as *Laminin B1 (LanB1)* and the tightly linked *viking* and *Collagen Type IV alpha1 (Col4a1)* genes, identified nine alleles of *LanB1* and one each of *viking* and, leaving eight complementation groups and six singletons.

### Whole genome sequencing

At least three alleles per complementation group, all singleton mutations, and the isogenic target chromosome were subjected to whole genome sequencing. For each allele, 10-15 late third instar larvae homozygous for the relevant mutagenized *M[3xP3-RFP.attP]\*\*\** chromosome were collected, washed, and frozen. Genomic DNA extractions was performed using the Qiagen DNeasy Blood & Tissue Kit and and Illumina Nextera Flex DNA Library Prep Kit was used to prepare indexed next-generation sequencing libraries following the manufacturer’s protocol. Genomic DNA was then provided to GTAC (Washington University) for whole genome sequencing at ≥30X coverage. Sequence reads were mapped to *Drosophila* reference BDGP6 using NovoAlign (Novocraft Technologies). Sentieon software was used to remove duplicate reads, realign reads around indels, recalibrate quality scores and call variants. The Sentieon DNAscope tool was used to identify variants. Joint variant calls were made using the mutant lines and a reference line. Variants were annotated using SnpEff and the variants were filtered using SnpSift to select unique coding variants within previously determined genomic intervals. Ultimately, this process identified nucleotide differences in coding regions and splice sites and small deletions that differed between the mutagenized chromosome and the isogenic target chromosome, which are likely EMS-induced mutations. For singletons, this approach identified a list of putative EMS-induced mutations, one of which is likely the causative lesion. For complementation groups, this approach identified the likely causative lesion for all complementation groups, as only one gene in each complementation group harbored independent EMS-induced mutations in its coding region in all sequenced alleles. When possible, we confirmed correspondence between gene and mutant phenotype via complementation crosses with appropriate deficiencies and gene-specific alleles and phenotypic analysis. All putative causative lesions were confirmed via PCR-based methods and Sanger sequencing (GeneWiz Inc.).

#### Genetic mapping and Complementation Crosses

To identify the causative lesion in singleton alleles, we took two approaches. For three lines, we used complementation crosses against deficiencies or gene-specific alleles that uncovered mutations likely to cause the associated CNS phenotype to confirm correspondence between gene and mutant phenotype. This approach identified mutations in *diaphanous*, *pvf3*, and *lethal giant larvae* as the causative to the CNS phenotype. For the remaining three singletons, we used the approach of Zhai et al (Zhai, Hiesinger et al. 2003) to map the causative lesion in each line on the genetic map relative to four P[w+] element inserts located at genetic map positions 13, 52, 62, and 86 in the second chromosome. Using this approach, we mapped one singleton to genetic map position 18-23 and 53-57 in the second chromosome; the results for the third singleton were inconclusive likely due to the presence of two or more lethal mutations in the chromosome. Complementation crosses against appropriate deficiencies and gene-specific alleles followed by phenotypic analysis identified *Tango1* as the singleton on 2L and *worniu* as the singleton near genetic map position 53-57. The two putative causative lesions were confirmed by PCR-based methods and Sanger sequencing.

#### Transgene construction

The *UAS-senseless-2* transgene was generated by amplifying nucleotides 276-2529 of the *sens-2 cDNA*, FI 20031 (accession number BT33368). The resulting PCR product was cloned into NotI and KpnI sites of pUAS-T. DNA was co-injected with helper plasmid into embryos of the genotype w1118 by Rainbow Transgenic Flies, Inc.

Due to the large size of *ppn*, the UAS-Ppn[RG] transgene was constructed piecemeal through PCR-based approaches amplified defined regions of partial cDNA clones and genomic DNA; these individual regions were then stitched together via Gibson cloning. First, we used the GH25513 cDNA clone, which contained a portion of the 5’ coding region of Ppn (accession number BT100225) to amplify nucleotides 1-2471 of the coding region of the Ppn[RG] transcript starting with the ATG start codon in exon 1 and continuing into exon 6. Second, we used genomic DNA from a strain of flies isogenic for a wild-type third chromosome to amplify nucleotides 2452-5205 within exon 6 nucleotide 5162-5384 (exons 7) of the Ppn[RG] transcript. Third, we used the partial 3’ GH05059 cDNA clone (accession number AY060635) to amplify in separate reactions nucleotides 5344-6602 (exons 8-9) and nucleotides 6556-8526 (exons 10 through 17, including *the* ATT stop-codon) of the Ppn[RG] transcript. The PCR products were assembled via Gibson cloning and inserted into the KpnI and XbaI sites of the phiC-based construct pJFRC28, which was then injected into the P[CaryP] attp2 fly line (BDSC: 8622) with phiC integrase by Rainbow Transgenic Flies, Inc.

#### Gene rescue and in vivo RNAi phenocopy assays

To restrict UAS-linked transgene expression specifically to glia, we used the *repoGAL4* driver line. To restrict UAS-linked transgene expression specifically to neurons, we paired the *elavGAL4* driver line, which activates transgene expression strongly in all neurons and moderately in glia, with *repoGAL80*, which blocks GAL4-dependent activation in glia (Awasaki, Lai et al. 2008). Gene rescue experiments were performed in the *sens-2-T2A-GAL4/sens-2^js4^* background.

#### Allele nomenclature

Alleles were given with their final designation (e.g., js1, js2) after they were placed in complementation groups. Within each complementation group alleles were named based on the order in which they identified in the screen. The one singleton mutation was designated *jsz4*.

#### Gene expression analysis

Gene expression analysis was performed essentially as described in (Patel 1994). Briefly, the larval CNS was dissected in PBS, fixed in 3.7% formaldehyde for 35 minutes, and washed in PTx (1xPBS, 0.1% TritonX-100) five times over one hour prior to primary antibody incubation. Fixed tissues was incubated in primary antibody with gentle rocking overnight at room temperature, then washed five times in PTx over the course of an hour, incubated in the appropriate secondary overnight for at least 4 hours to overnight at room temperature, and then washed again in PTx at least five times over one hour. The CNS was then dissected and mounted in PTx or in DPX mountant after dehydration via an ethanol series and clearing in xylenes (Truman, Talbot et al. 1994). All imaging was performed on a Zeiss LSM-700 Confocal Microscope, using Zen software. The following antibodies were used in the study: Deadpan (1:2000, (Skeath, Wilson et al. 2017)); rat monoclonal antibody 7E8A10 (ELAV; 1:200; (O’Neill, Rebay et al. 1994)); mouse monoclonal antibody 8D12 (REPO; 1:100) (Alfonso and Jones 2002); rabbit anti-Sens-2 (this paper; 1:500); rabbit anti-Ppn (this paper; 1:250); goat Anti-GFP Dylight TM 488 Conjugated, preadsorbed, (Rockland; #600-141-215). Secondary antibodies were obtained from the following sources. The Alexa Fluor Plus 488 secondary antibodies of appropriate species specificity were obtained from ThermoFisher and used at a dilution of 1:400: e.g., Donkey anti-Rat IgG Alexa Fluor™ Plus 488; catalog number Cat # A-21208. The Cy5 secondary antibodies of appropriate species specificity were obtained from Jackson Immunoresearch and used at a dilution of 1:400: e.g., Cy5 Donkey Anti-Rat IgG, #712-175-153.

#### Antibody generation

To generate the *Senseless-2* antibody, DNA encoding amino acids 1-199 of Senseless-2 was inserted into pET-29a (Novagen) for protein expression and purification. Protein-specific antibody responses were mounted in rabbits (Pocono Rabbit Farm and Laboratory, PA, USA) and the resulting sera was used at a 1:500-1:1000 dilution. As all senseless-2 mutations arise C-terminal to the epitopes against which the antibody was generated, we confirmed specificity of this antibody to senseless-2 by driving UAS-senseless-2 expression in subperineural glia within the ventral nerve cord, which normally do not express *senseless-2*, and detecting high levels of Senseless-2 protein in these cells (Fig. S1).

#### Papilin antibody generation

YenZym (CA, USA) was used as a commercial source to generate affinity purified antibodies against two distinct synthetic peptides that corresponded to amino acids 27-47 (RFPGLRQKRQYGANMYLPEC) and 2670-2691 (TRPVTQRPSYPYRPTRPAYVPE) of Ppn-PG. Briefly, each peptide was conjugated to KLH, used as an immunogen in rabbits to generate a peptide-specific antibody response, and antibodies specific to the peptide were affinity purified. The two affinity purified antibodies were combined and used at a dilution of 1:250-1:500 for immunofluorescence analysis.

#### Generation of senseless-2 CRIMIC T2A-GAL4 line

To generate sens-2-GAL4-DBD, we used a modified version of the strategy developed by Kanca et al. (Kanca, Zirin et al. 2019). We used the Genewiz company to synthesize a DNA fragment into the EcoRV site of the pUC-GW-Kan vector. This fragment is made of the left and right homology arms (HA) which are immediately adjacent to the cut site and restriction enzyme sites (SacI-KpnI) between these arms. We then directionally cloned the attp-FRT-splitGAL4-FRT-attp fragment (see below) into the middle of left and right HAs using the SacI and KpnI sites. We used the genome sequence of the injected flies (vasa-cas9, BDSC-51324) for design.

### HA flanking sequences

Left HA: 5’cgtgtgtgagagaga 3’aacggtcttttccct; Right HA: 5’gggtggggcagcgcc 3’tatctatcatagata

### SacI-attp-FRT-splitGAL4-FRT-attp-Kpn1 fragment

We used the Gibson cloning method to clone three fragments into pBS-KS digested with SacI and KpnI. Primer pairs and templates shown below: Note we also included ECoRI and EcoRV sites for replacing the trojan exon between attpFRT sites as a back-up strategy when needed.

1. *SacI-*attp-FRT from pM14 (Kanca, Zirin et al. 2019):

a. F <u>actcactatagggcgaattgGAGCTC</u>*acggacacaccgaag*
b. R *caagtcgccatgttggatcgac*
2. Split-GAL4 from Trojan-AD/DBD (Diao, Ironfield et al. 2015):

a. F cta*gaaagtataggaacttc*GAATTC**agtcgatccaacatggcgacttg**
b. R cttt*ctagagaataggaacttc*GATATC**aaacgagtttttaagcaaactcactcc**
3. Kpn1-attp-FRT from pM14 (Kanca, Zirin et al. 2019):

a. F **ggagtgagtttgcttaaaaactcgttt**GATATCgaag*ttcctattctctagaaag*
b. R <u>cactaaagggaacaaaagctgGGTACC</u>gtac*tgacggacacaccgaag*

Corresponding sequences from pBS-KS are underlined, pM14 are in italics, and Trojan AD/DBD are in bold; restriction enzyme sites are in all caps.

Guide RNAs were identified with CRISPR target Finder (Gratz, Ukken et al. 2014) with maximum stringency and minimal off-target effects. We used pCFD4 vector (Port, Chen et al. 2014) to clone sens-2 specific guide and donor linearizing guide RNAs via a PCR intermediate with the following primers:

*sens-2 guide RNA*: attttaacttgctatttctagctctaaaacCCCAACGGTCTTTTCCCTGC gacgttaaattgaaaataggtc *linearizing guide RNA*: tatataggaaagatatccgggtgaacttcGTAGTACGATCATAACAACG gttttagagctagaaatagcaag

All constructs were injected into *vasa-cas9* (BDSC #51324), which were then crossed to *Tubulin-GAL4-AD, UAS-TdTomato/CyO* and scored for TdTomato expression to identify positive lines. Verification of targeting was confirmed via PCR-based sequencing methods. After we generated the *sens-2-GAL4-DBD* line, we used the strategy described in Diao et al., (Diao, Ironfield et al. 2015) to replace the split-GAL4 insert with a T2A-GAL4 and then conducted all experiments with the sens-2-T2A-GAL4 line, termed *sens-2-GAL4*.

The *papilin*-T2A-GAL4 line was generated from the *ppn^MI03189^*insert following the genetic cross scheme outlined in Diao et al., 2015 (Diao, Ironfield et al. 2015).

### Fly Stocks used

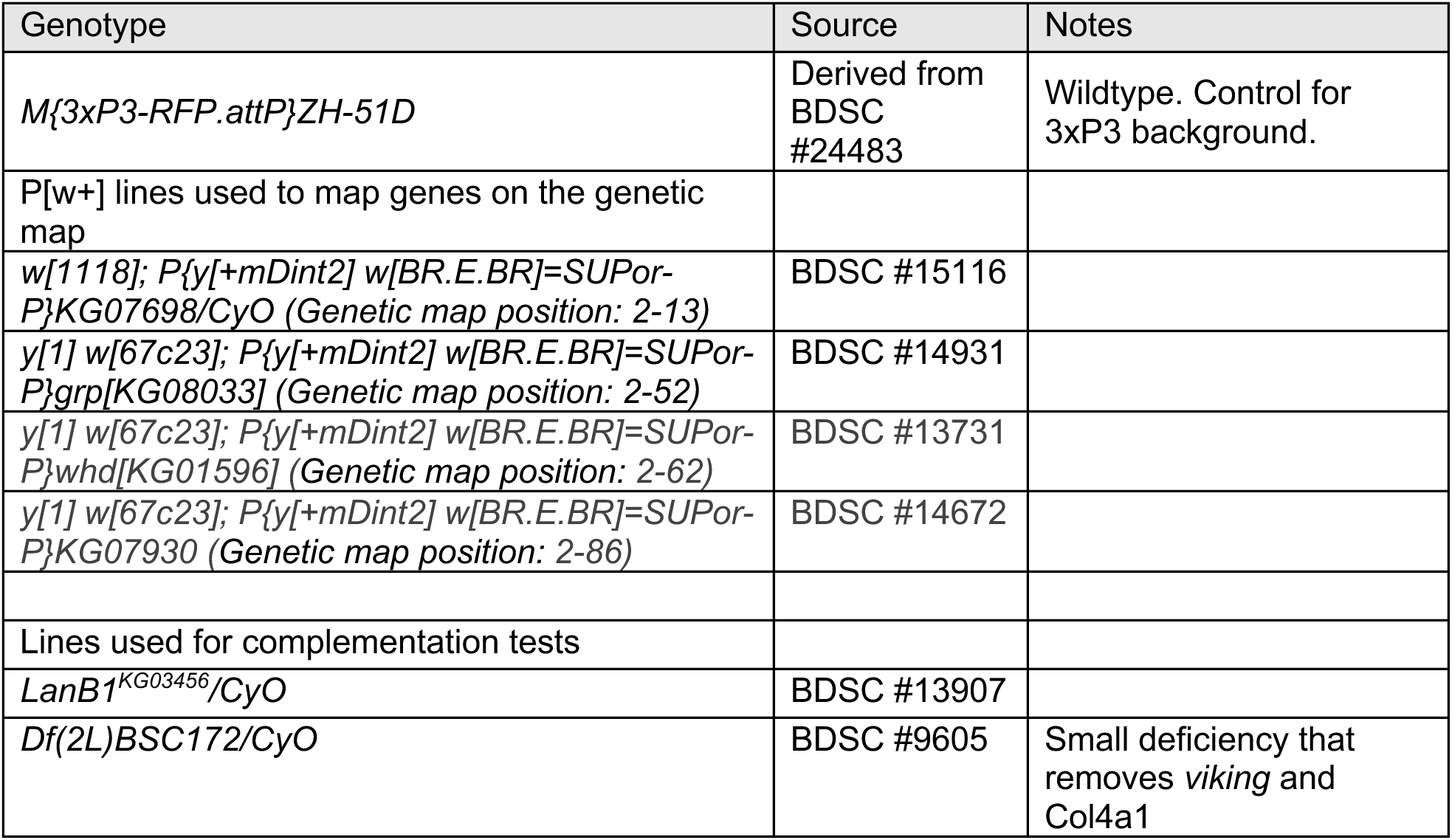
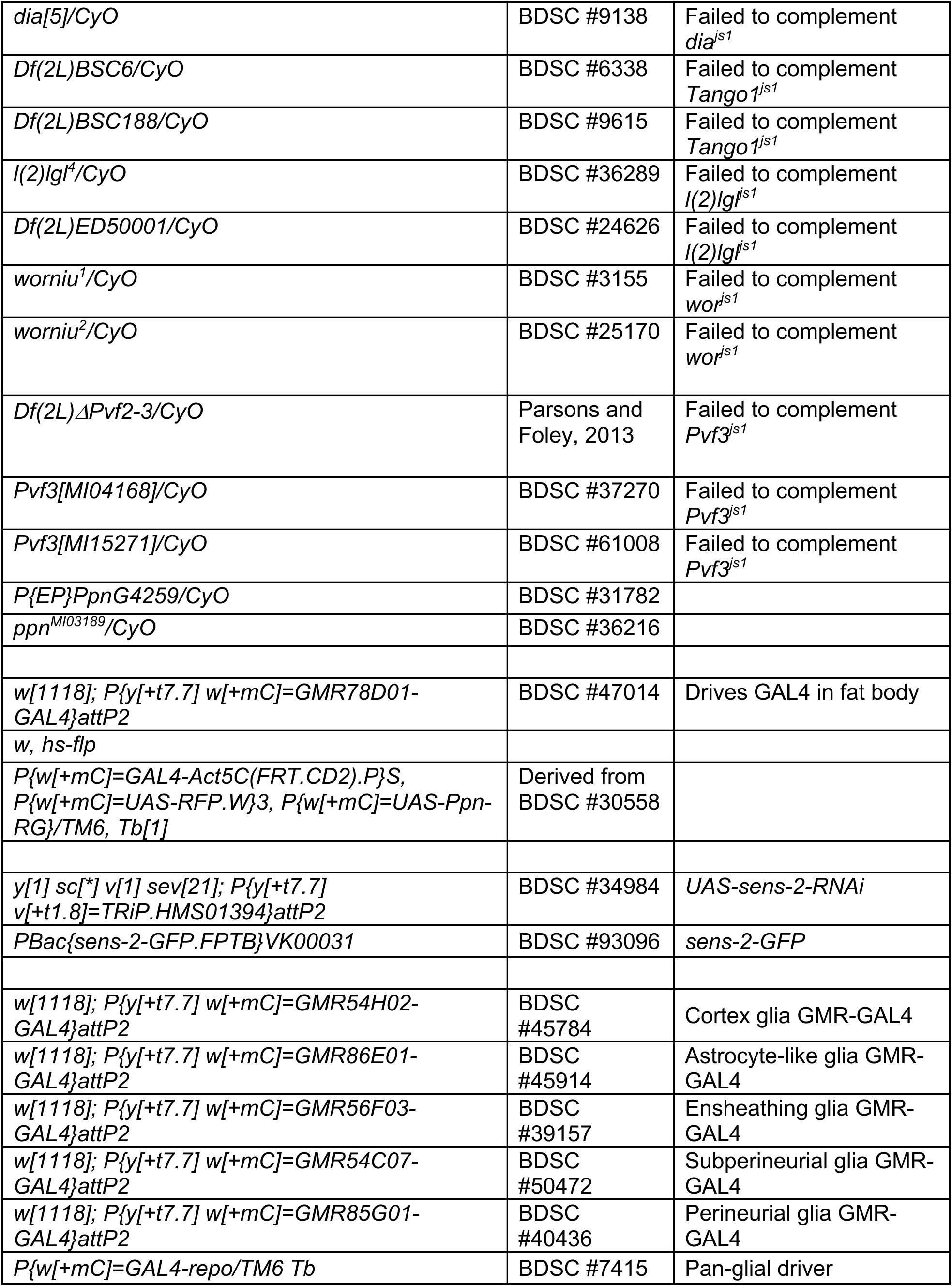
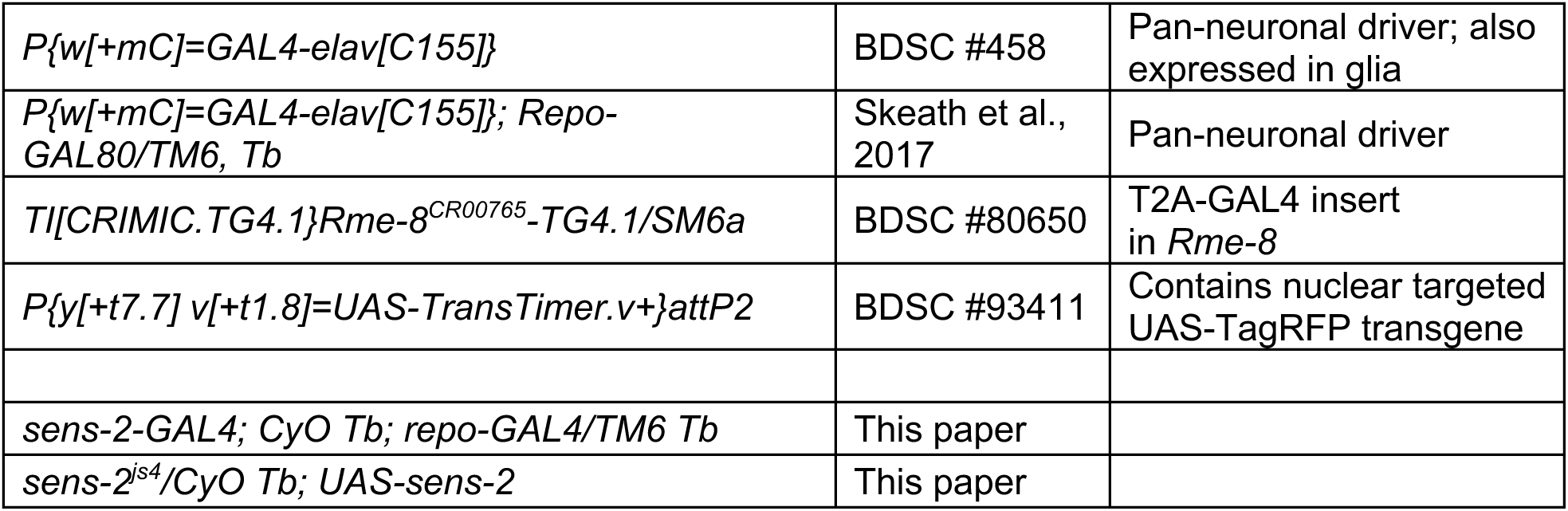

## Data Availability Statement

All data necessary for confirming the conclusions of this manuscript are represented within the article, its tables, figures, and supplementary information. All fly stocks are available upon request.

## Acknowledgements

We thank the Iowa Developmental Studies Hybridoma Bank for antibodies, and the Bloomington Stock Center for fly stocks. We thank the Genome Technology Access Center (GTAC) at Washington University for next-generation sequencing.

## References

Alfonso, T. B. and B. W. Jones (2002). “gcm2 promotes glial cell differentiation and is required with glial cells missing for macrophage development in Drosophila.” Dev Biol 248(2): 369–383.

Apte, S. S. (2009). “A disintegrin-like and metalloprotease (reprolysin-type) with thrombospondin type 1 motif (ADAMTS) superfamily: functions and mechanisms.” J Biol Chem 284(46): 31493–31497.

Ashraf, S. I., A. Ganguly, J. Roote and Y. T. Ip (2004). “Worniu, a Snail family zinc-finger protein, is required for brain development in Drosophila.” Dev Dyn 231(2): 379–386.

Awasaki, T., S. L. Lai, K. Ito and T. Lee (2008). “Organization and postembryonic development of glial cells in the adult central brain of Drosophila.” J Neurosci 28(51): 13742–13753.

Berger, C., S. Renner, K. Luer and G. M. Technau (2007). “The commonly used marker ELAV is transiently expressed in neuroblasts and glial cells in the Drosophila embryonic CNS.” Dev Dyn 236(12): 3562–3568.

Bigotti, M. G. and A. Brancaccio (2021). “High degree of conservation of the enzymes synthesizing the laminin-binding glycoepitope of alpha-dystroglycan.” Open Biol 11(9): 210104.

Campbell, A. G., L. I. Fessler, T. Salo and J. H. Fessler (1987). “Papilin: a Drosophila proteoglycan-like sulfated glycoprotein from basement membranes.” J Biol Chem 262(36): 17605–17612.

Castrillon, D. H. and S. A. Wasserman (1994). “Diaphanous is required for cytokinesis in Drosophila and shares domains of similarity with the products of the limb deformity gene.” Development 120(12): 3367–3377.

Dai, J., B. Estrada, S. Jacobs, B. J. Sanchez-Sanchez, J. Tang, M. Ma, P. Magadan-Corpas, J. C. Pastor-Pareja and M. D. Martin-Bermudo (2018). “Dissection of Nidogen function in Drosophila reveals tissue-specific mechanisms of basement membrane assembly.” PLoS Genet 14(9): e1007483.

Diao, F., H. Ironfield, H. Luan, F. Diao, W. C. Shropshire, J. Ewer, E. Marr, C. J. Potter, M. Landgraf and B. H. White (2015). “Plug-and-play genetic access to drosophila cell types using exchangeable exon cassettes.” Cell Rep 10(8): 1410–1421.

Gateff, E. (1994). “Tumor-suppressor genes, hematopoietic malignancies and other hematopoietic disorders of Drosophila melanogaster.” Ann N Y Acad Sci 712: 260–279.

Gateff, E. and H. A. Schneiderman (1974). “Developmental capacities of benign and malignant neoplasms ofDrosophila.” Wilhelm Roux Arch Entwickl Mech Org 176(1): 23–65.

Gomez-Lamarca, M., L. A. Snowdon, E. Seib, T. Klein and S. Bray (2015). “Rme-8 depletion perturbs Notch recycling and predisposes to pathogenic signaling.” J Cell Biol 210(3): 517.

Gratz, S. J., F. P. Ukken, C. D. Rubinstein, G. Thiede, L. K. Donohue, A. M. Cummings and K. M. O’Connor-Giles (2014). “Highly specific and efficient CRISPR/Cas9-catalyzed homology-directed repair in Drosophila.” Genetics 196(4): 961–971.

Hannun, Y. A. and L. M. Obeid (2018). “Sphingolipids and their metabolism in physiology and disease.” Nat Rev Mol Cell Biol 19(3): 175–191.

He, L., R. Binari, J. Huang, J. Falo-Sanjuan and N. Perrimon (2019). “In vivo study of gene expression with an enhanced dual-color fluorescent transcriptional timer.” Elife 8.

Hermanstyne, T. O., L. Johnson, K. M. Wylie and J. B. Skeath (2022). “Helping others enhances graduate student wellness and mental health.” Nat Biotechnol 40(4): 618–619.

Isabella, A. J. and S. Horne-Badovinac (2015). “Building from the Ground up: Basement Membranes in Drosophila Development.” Curr Top Membr 76: 305–336.

Jung, W. H., C. C. Liu, Y. L. Yu, Y. C. Chang, W. Y. Lien, H. C. Chao, S. Y. Huang, C. H. Kuo, H. C. Ho and C. C. Chan (2017). “Lipophagy prevents activity-dependent neurodegeneration due to dihydroceramide accumulation in vivo.” EMBO Rep 18(7): 1150–1165.

Kanca, O., J. Zirin, J. Garcia-Marques, S. M. Knight, D. Yang-Zhou, G. Amador, H. Chung, Z. Zuo, L. Ma, Y. He, W. W. Lin, Y. Fang, M. Ge, S. Yamamoto, K. L. Schulze, Y. Hu, A. C. Spradling, S. E. Mohr, N. Perrimon and H. J. Bellen (2019). “An efficient CRISPR-based strategy to insert small and large fragments of DNA using short homology arms.” Elife 8.

Karkali, K., P. Tiwari, A. Singh, S. Tlili, I. Jorba, D. Navajas, J. J. Munoz, T. E. Saunders and E. Martin-Blanco (2022). “Condensation of the Drosophila nerve cord is oscillatory and depends on coordinated mechanical interactions.” Dev Cell 57(7): 867–882 e865.

Keeley, D. P., E. Hastie, R. Jayadev, L. C. Kelley, Q. Chi, S. G. Payne, J. L. Jeger, B. D. Hoffman and D. R. Sherwood (2020). “Comprehensive Endogenous Tagging of Basement Membrane Components Reveals Dynamic Movement within the Matrix Scaffolding.” Dev Cell 54(1): 60–74 e67.

Kim, S. N., A. Jeibmann, K. Halama, H. T. Witte, M. Walte, T. Matzat, H. Schillers, C. Faber, V. Senner, W. Paulus and C. Klambt (2014). “ECM stiffness regulates glial migration in Drosophila and mammalian glioma models.” Development 141(16): 3233–3242.

Kramerova, I. A., N. Kawaguchi, L. I. Fessler, R. E. Nelson, Y. Chen, A. A. Kramerov, M. Kusche-Gullberg, J. M. Kramer, B. D. Ackley, A. L. Sieron, D. J. Prockop and J. H. Fessler (2000). “Papilin in development; a pericellular protein with a homology to the ADAMTS metalloproteinases.” Development 127(24): 5475–5485.

Kudron, M. M., A. Victorsen, L. Gevirtzman, L. W. Hillier, W. W. Fisher, D. Vafeados, M. Kirkey, A. S. Hammonds, J. Gersch, H. Ammouri, M. L. Wall, J. Moran, D. Steffen, M. Szynkarek, S. Seabrook-Sturgis, N. Jameel, M. Kadaba, J. Patton, R. Terrell, M. Corson, T. J. Durham, S. Park, S. Samanta, M. Han, J. Xu, K. K. Yan, S. E. Celniker, K. P. White, L. Ma, M. Gerstein, V. Reinke and R. H. Waterston (2018). “The ModERN Resource: Genome-Wide Binding Profiles for Hundreds of Drosophila and Caenorhabditis elegans Transcription Factors.” Genetics 208(3): 937–949.

Lee, P. T., J. Zirin, O. Kanca, W. W. Lin, K. L. Schulze, D. Li-Kroeger, R. Tao, C. Devereaux, Y. Hu, V. Chung, Y. Fang, Y. He, H. Pan, M. Ge, Z. Zuo, B. E. Housden, S. E. Mohr, S. Yamamoto, R. W. Levis, A. C. Spradling, N. Perrimon and H. J. Bellen (2018). “A gene-specific T2A-GAL4 library for Drosophila.” Elife 7.

Li, G., M. G. Forero, J. S. Wentzell, I. Durmus, R. Wolf, N. C. Anthoney, M. Parker, R. Jiang, J. Hasenauer, N. J. Strausfeld, M. Heisenberg and A. Hidalgo (2020). “A Toll-receptor map underlies structural brain plasticity.” Elife 9.

Lin, Y. R., B. V. Reddy and K. D. Irvine (2008). “Requirement for a core 1 galactosyltransferase in the Drosophila nervous system.” Dev Dyn 237(12): 3703–3714.

Malhotra, V. and P. Erlmann (2011). “Protein export at the ER: loading big collagens into COPII carriers.” EMBO J 30(17): 3475–3480.

Meyer, S., I. Schmidt and C. Klambt (2014). “Glia ECM interactions are required to shape the Drosophila nervous system.” Mech Dev 133: 105–116.

Miller, C. M., N. Liu, A. Page-McCaw and H. T. Broihier (2011). “Drosophila MMP2 regulates the matrix molecule faulty attraction (Frac) to promote motor axon targeting in Drosophila.” J Neurosci 31(14): 5335–5347.

Morin, X., R. Daneman, M. Zavortink and W. Chia (2001). “A protein trap strategy to detect GFP-tagged proteins expressed from their endogenous loci in Drosophila.” Proc Natl Acad Sci U S A 98(26): 15050–15055.

Norris, A., C. T. McManus, S. Wang, R. Ying and B. D. Grant (2022). “Mutagenesis and structural modeling implicate RME-8 IWN domains as conformational control points.” PLoS Genet 18(10): e1010296.

O’Neill, E. M., I. Rebay, R. Tjian and G. M. Rubin (1994). “The activities of two Ets-related transcription factors required for Drosophila eye development are modulated by the Ras/MAPK pathway.” Cell 78(1): 137–147.

Olofsson, B. and D. T. Page (2005). “Condensation of the central nervous system in embryonic Drosophila is inhibited by blocking hemocyte migration or neural activity.” Dev Biol 279(1): 233–243.

Page-McCaw, A., J. Serano, J. M. Sante and G. M. Rubin (2003). “Drosophila matrix metalloproteinases are required for tissue remodeling, but not embryonic development.” Dev Cell 4(1): 95–106.

Parsons, B. and E. Foley (2013). “The Drosophila platelet-derived growth factor and vascular endothelial growth factor-receptor related (Pvr) protein ligands Pvf2 and Pvf3 control hemocyte viability and invasive migration.” J Biol Chem 288(28): 20173–20183.

Pastor-Pareja, J. C. and T. Xu (2011). “Shaping cells and organs in Drosophila by opposing roles of fat body-secreted Collagen IV and perlecan.” Dev Cell 21(2): 245–256.

Patel, N. H. (1994). “Imaging neuronal subsets and other cell types in whole-mount Drosophila embryos and larvae using antibody probes.” Methods Cell Biol 44: 445–487.

Petley-Ragan, L. M., E. L. Ardiel, C. H. Rankin and V. J. Auld (2016). “Accumulation of Laminin Monomers in Drosophila Glia Leads to Glial Endoplasmic Reticulum Stress and Disrupted Larval Locomotion.” J Neurosci 36(4): 1151–1164.

Port, F., H. M. Chen, T. Lee and S. L. Bullock (2014). “Optimized CRISPR/Cas tools for efficient germline and somatic genome engineering in Drosophila.” Proc Natl Acad Sci U S A 111(29): E2967–2976.

Praissman, J. L., D. H. Live, S. Wang, A. Ramiah, Z. S. Chinoy, G. J. Boons, K. W. Moremen and L. Wells (2014). “B4GAT1 is the priming enzyme for the LARGE-dependent functional glycosylation of alpha-dystroglycan.” Elife 3.

Rodriguez, A., H. Oliver, H. Zou, P. Chen, X. Wang and J. M. Abrams (1999). “Dark is a Drosophila homologue of Apaf-1/CED-4 and functions in an evolutionarily conserved death pathway.” Nat Cell Biol 1(5): 272–279.

Rodriguez-Manzaneque, J. C., R. Fernandez-Rodriguez, F. J. Rodriguez-Baena and M. L. Iruela-Arispe (2015). “ADAMTS proteases in vascular biology.” Matrix Biol 44-46:38-45.

Saito, K., M. Chen, F. Bard, S. Chen, H. Zhou, D. Woodley, R. Polischuk, R. Schekman and V. Malhotra (2009). “TANGO1 facilitates cargo loading at endoplasmic reticulum exit sites.” Cell 136(5): 891–902.

Skeath, J. B., B. A. Wilson, S. E. Romero, M. J. Snee, Y. Zhu and H. Lacin (2017). “The extracellular metalloprotease AdamTS-A anchors neural lineages in place within and preserves the architecture of the central nervous system.” Development 144(17): 3102–3113.

Stork, T., D. Engelen, A. Krudewig, M. Silies, R. J. Bainton and C. Klambt (2008). “Organization and function of the blood-brain barrier in Drosophila.” J Neurosci 28(3): 587–597.

Truman, J. W., W. S. Talbot, S. E. Fahrbach and D. S. Hogness (1994). “Ecdysone receptor expression in the CNS correlates with stage-specific responses to ecdysteroids during Drosophila and Manduca development.” Development 120(1): 219–234.

Wilson, D. G., K. Phamluong, L. Li, M. Sun, T. C. Cao, P. S. Liu, Z. Modrusan, W. N. Sandoval, L. Rangell, R. A. Carano, A. S. Peterson and M. J. Solloway (2011). “Global defects in collagen secretion in a Mia3/TANGO1 knockout mouse.” J Cell Biol 193(5): 935–951.

Yildirim, K., J. Petri, R. Kottmeier and C. Klambt (2019). “Drosophila glia: Few cell types and many conserved functions.” Glia 67(1): 5–26.

Yoshida-Moriguchi, T., L. Yu, S. H. Stalnaker, S. Davis, S. Kunz, M. Madson, M. B. Oldstone, H. Schachter, L. Wells and K. P. Campbell (2010). “O-mannosyl phosphorylation of alpha-dystroglycan is required for laminin binding.” Science 327(5961): 88–92.

Yurchenco, P. D. (2011). “Basement membranes: cell scaffoldings and signaling platforms.” Cold Spring Harb Perspect Biol 3(2).

Zhai, R. G., P. R. Hiesinger, T. W. Koh, P. Verstreken, K. L. Schulze, Y. Cao, H. Jafar-Nejad, K. K. Norga, H. Pan, V. Bayat, M. P. Greenbaum and H. J. Bellen (2003). “Mapping Drosophila mutations with molecularly defined P element insertions.” Proc Natl Acad Sci U S A 100(19): 10860–10865.

Zhang, Y., B. Grant and D. Hirsh (2001). “RME-8, a conserved J-domain protein, is required for endocytosis in Caenorhabditis elegans.” Mol Biol Cell 12(7): 2011–2021.

Zhu, Y., K. Cho, H. Lacin, Y. Zhu, J. DiPaola, B. A. Wilson, G. J. Patti and J. B. Skeath (2024). “Dihydroceramide desaturase governs endoplasmic reticulum and lipid droplet homeostasis to promote glial function in the nervous system.” BioRxiv.

