## Supplemental Figures and Figure legends for "A Genetic Screen in *Drosophila* uncovers a role for *senseless-2* in surface glia in the peripheral nervous system to regulate CNS morphology"

**Figure S1)**

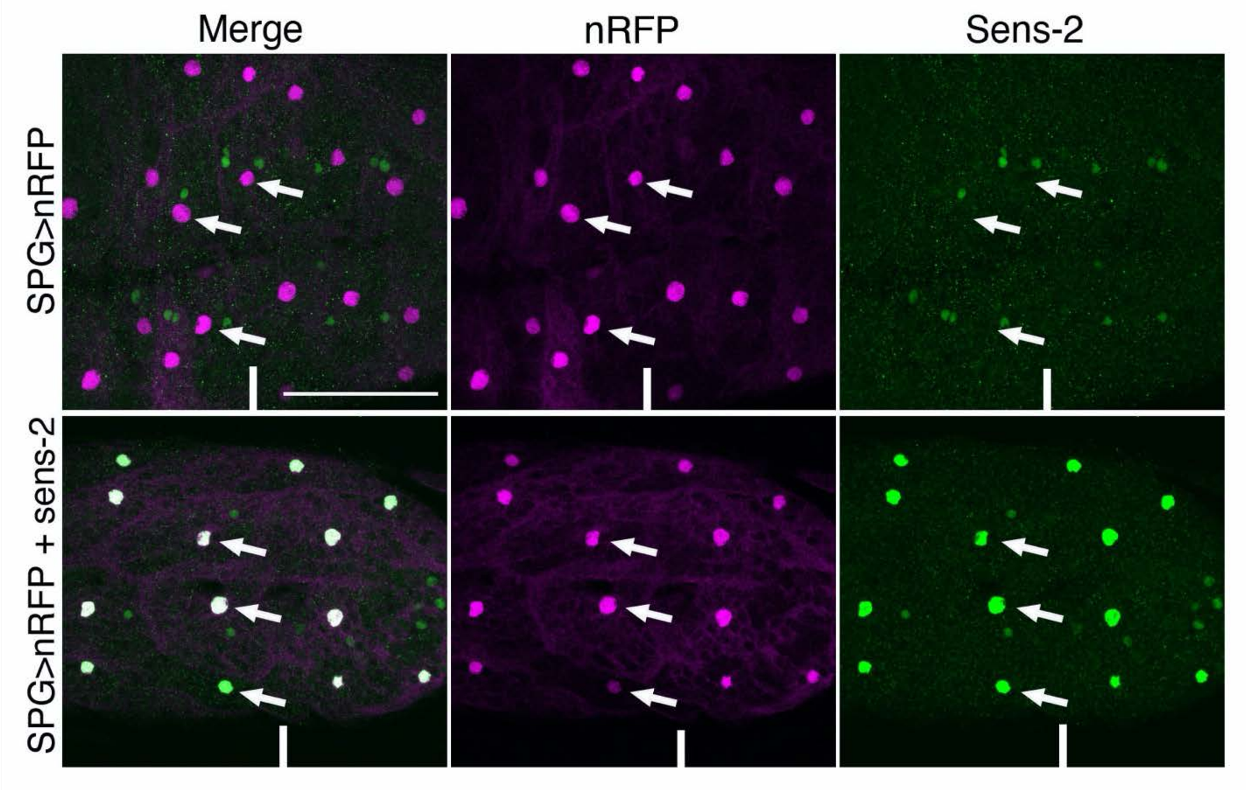


**Figure S1)** Senseless-2 antibody recognizes ectopically expressed Senseless-2 protein. High magnification views of the abdominal region of the ventral nerve cord from late third instar larvae expressing either nuclear-RFP (nRFP; top) or nRFP and senseless-2 (bottom) under the control of the subperineural glia specific GAL4 line GMR54C07-GAL4. Top: The senseless-2 antibody does not detect senseless protein in subperineural glia in otherwise wild-type larvae that express nRFP in subperineural glia (arrows). Bottom: The senseless-2 antibody strongly detects Senseless-2 protein in subperineurial glia upon GAL4-mediated expression of senseless-2 in these cells (arrows). Line represents midline of ventral nerve cord. Anterior is up; scale bar is 50 microns.

Figure S2)

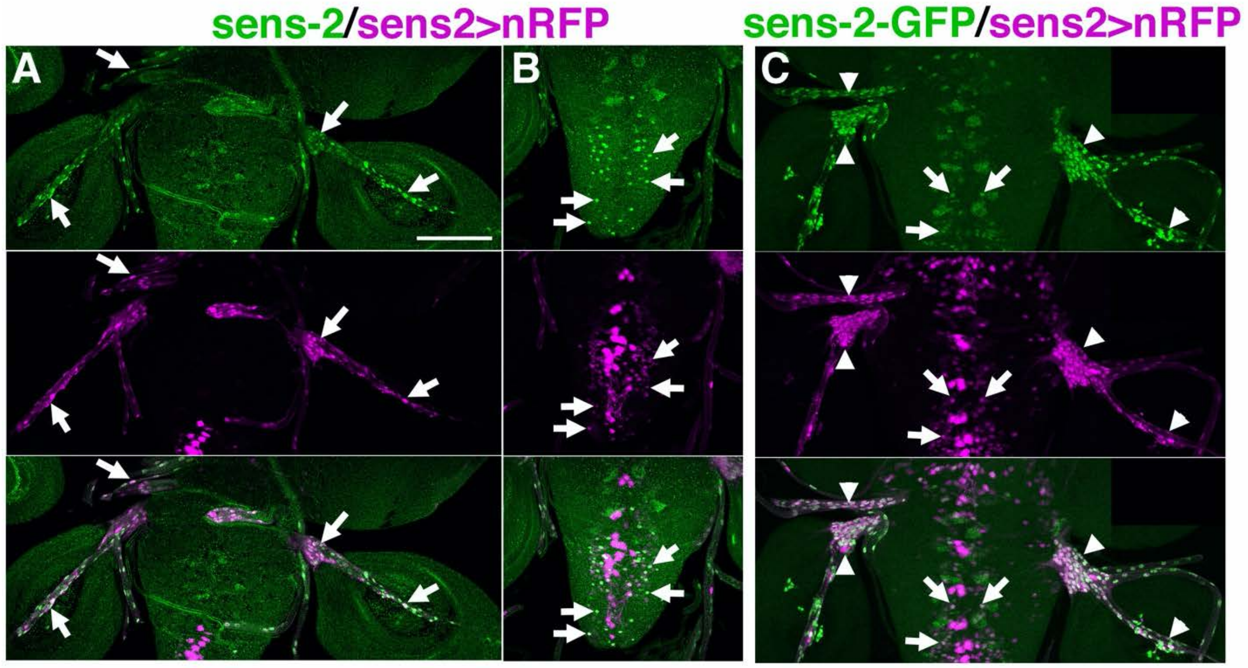


**Figure S2)** Coincident expression of sens-2 protein, sens-2-T2A-GAL4 CRIMIC line, and sens-2-GFP line.

A-B) Ventral views of CNS and peripheral nerves of late third instar *sens-2-T2A-GAL4>nRFP* larvae labeled with sens-2 antibody (green) and nRFP (magenta) highlighting *sens-2* expression in peripheral nerves (A) and within the CNS (B). Arrows in A highlight co-expression of Sens-2 and nRFP in peripheral glia; arrows in B highlight co-expression of sens-2 and nRFP in neurons. C) Ventral views of CNS and peripheral nerves of late third instar *sens-2-T2A-GAL4>nRFP* marked for GFP and nRFP. Arrows highlight co-expression in peripheral glia; arrowheads identify co-expression in neurons. Anterior is up; scale bar is 100 µm.

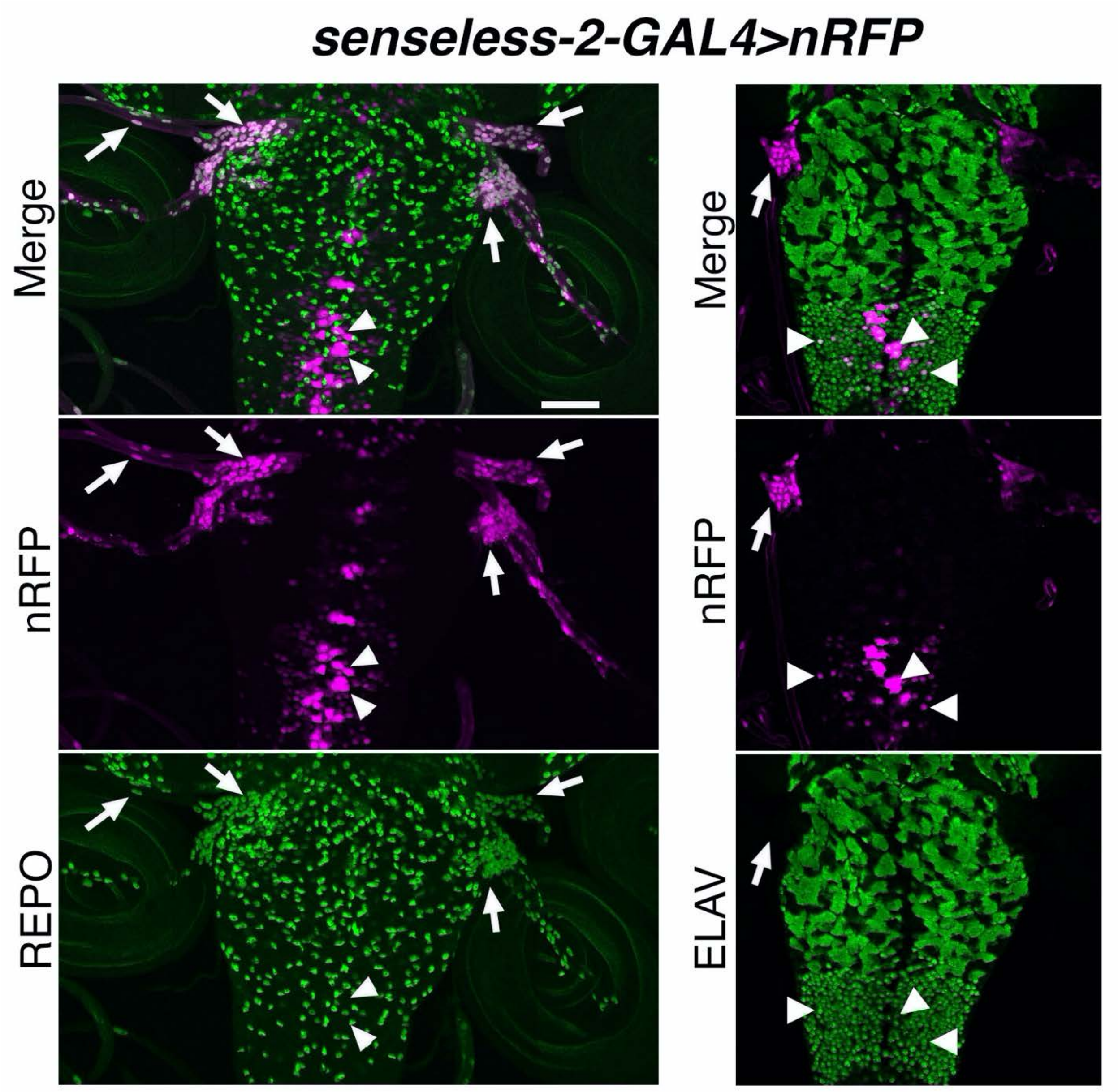


Figure S3)

Figure S3) *sens-2-GAL4* labels neurons in the CNS and peripheral glia. Ventral views of the CNS and peripheral nerves of late third instar *sens-2-GAL4>nRFP* larvae labeled for RFP and repo to mark glia (green; left) or ELAV to mark neurons (green; right). Left) Expression of nRFP (magenta) is co-expressed with REPO (green) in peripheral glia, but not in any cells in the CNS. Right) Expression of nRFP (magenta) is co-expressed with ELAV (green) in neurons in the CNS (arrowheads). Arrows point to peripheral glia; arrowheads to sens-2-positive neurons. Anterior is up; scale bar is 100 µm.
